# Pathological changes induced by Alzheimer’s brain inoculation in amyloid-beta plaque-bearing mice

**DOI:** 10.1101/2021.04.06.438654

**Authors:** Suzanne Lam, Anne-Sophie Hérard, Susana Boluda, Fanny Petit, Sabiha Eddarkaoui, Karine Cambon, The Brainbank Neuro-CEB Neuropathology Network, Jean-Luc Picq, Luc Buée, Charles Duyckaerts, Stéphane Haïk, Marc Dhenain

## Abstract

Alzheimer’s disease (AD) is characterized by intracerebral accumulations of extracellular amyloid-β (Aβ) plaques and intracellular tau pathology that spread in the brain. Tau lesions occur in the form of neuropil threads, neurofibrillary tangles, and neuritic plaques *i.e.* tau aggregates within neurites surrounding Aβ deposits. The cascade of events linking these lesions and synaptic or memory impairments are still debated. Intracerebral infusion of human AD brain extracts in Aβ plaque-bearing mice that do not overexpress pathological tau proteins induces tau pathologies following heterotopic seeding of mouse tau protein. There is however little information regarding the downstream events including synaptic or cognitive repercussions of tau pathology induction in these models. In the current study, human AD brain extracts (AD_be_) and control-brain extracts (Ctrl_be_) were infused in the hippocampus of Aβ plaque-bearing APP_swe_/PS1_dE9_ mice. Memory, synaptic density, as well as Aβ plaque and tau aggregate loads, microgliosis, astrogliosis at the inoculation site and in connected regions (perirhinal/entorhinal cortex) were evaluated 4 and 8 months post-inoculation. AD_be_ inoculation induced memory deficit. It increased deposition of Aβ plaques close to the inoculation site. Tau pathology was also induced in AD_be_-inoculated mice. Neuropil threads and neurofibrillary tangles occurred next to the inoculation site and spread to connected regions notably the perirhinal/entorhinal cortex. Neuritic plaque pathology was detected in both AD_be_- and Ctrl_be_- inoculated animals but AD_be_ inoculation increased the severity close and at distance of the inoculation site. Finally, AD_be_ inoculation reduced synaptic density close to the inoculation site and in connected regions as the perirhinal/entorhinal cortex. Synaptic impairments were correlated with increased severity of neuritic plaques but not of other tau lesions or Aβ lesions, which suggests that neuritic plaques are a culprit for synaptic loss. Synaptic density was also associated with microglial load.

**Graphical abstract:** 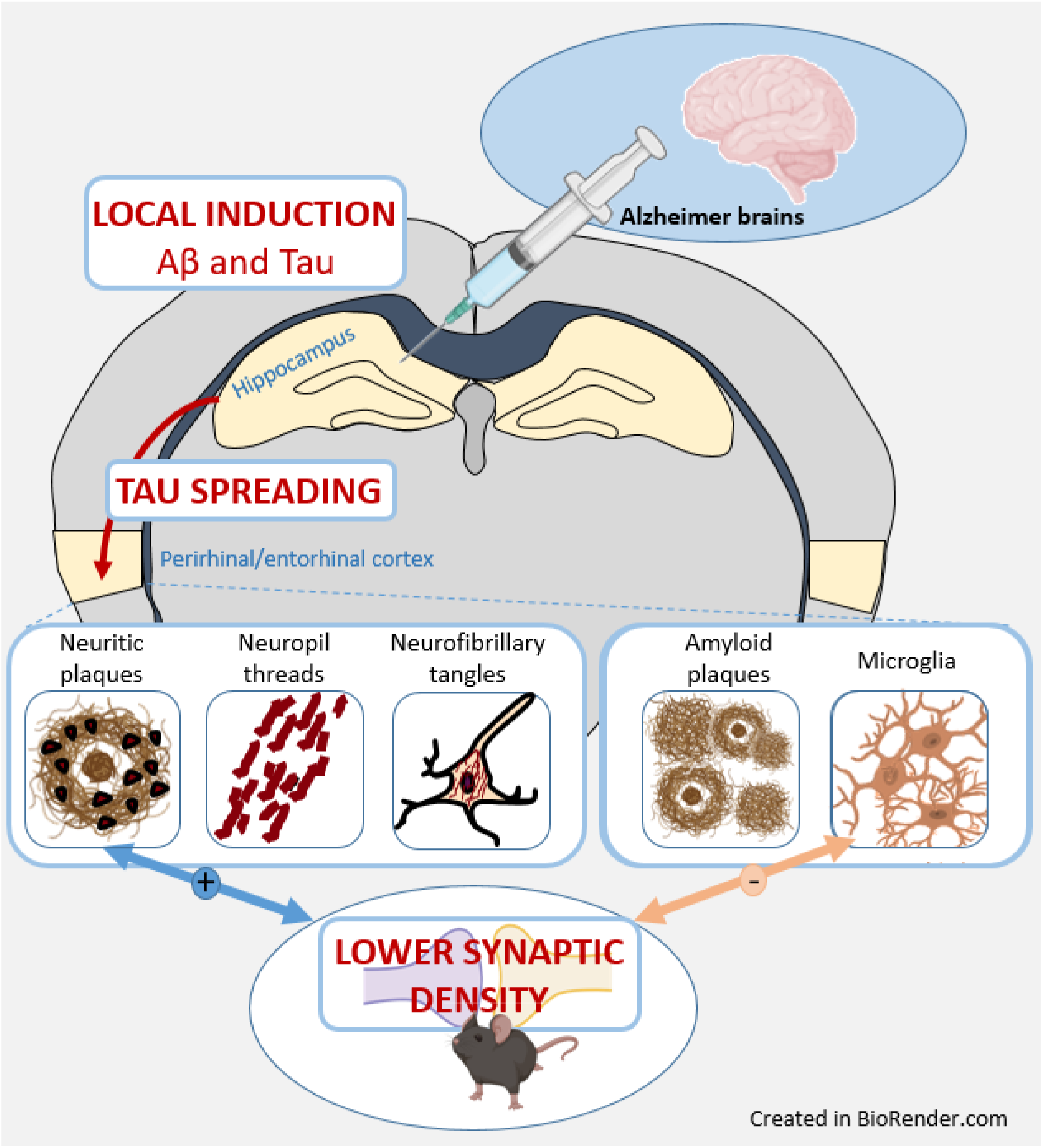

## Introduction

Alzheimer’s disease (AD) is a clinical entity leading to cognitive deficits. AD core lesions include amyloid-β (Aβ) plaques and intracellular tau accumulations that spread in a highly stereotyped pattern through the brain [3]. In humans, during AD, memory impairment is correlated with neocortical tau pathology and with synaptic alterations but not as well with Aβ plaque load [2, 18, 30]. Transgenic animals are widely used to investigate pathophysiological mechanisms associated with AD. Association between synaptic deficits and Aβ pathology was reported in transgenic (Tg) mouse models overexpressing Aβ [26–28] while association between synaptic deficits and tau pathology was outlined in Tg models overexpressing tau protein (with or without frontotemporal dementia mutations) [7, 14, 22, 28]. These results suggest a role of both Aβ and tau pathologies in the induction of synaptic deficits in rodents, but the relative participation of Aβ and tau lesions to synaptic impairments remains undefined. As AD is characterized by both Aβ and tau pathologies, it is critical to investigate its pathophysiology in animal models presenting with both Aβ and tau lesions. Mouse models of amyloidosis were shown to display tau lesions mainly in neurites close to Aβ deposits [17, 19, 23]. Presumably due to the low level of tau pathology, these animals are not used as models of both Aβ and tau pathologies. Novel categories of models with both Aβ and tau pathologies were recently established by infusing human AD brain extracts (AD_be_) in the brain of Aβ plaque-bearing mice that do not overexpress pathological tau proteins [13, 31]. These models rely on the notion that tau proteins have prion-like properties and can be experimentally transmitted. Once in the cell environment, they can be taken up by unaffected cells, and are amplified by seeding the aggregation of endogenous tau. In these models, tau seeds from AD_be_ induce the heterotopic seeding of mouse tau [1, 31]. These models were mainly used to investigate the relationships between Aβ and tau pathologies [13, 16, 31]. The relationships between AD core lesions and downstream events as synaptic impairments has not been reported, and memory changes remain mostly undefined in these models [13, 31]. To address these questions, we infused AD_be_ as well as control brain extracts (Ctrl_be_) in the hippocampus of APP_swe_/PS1_dE9_ mice that have a high Aβ production, express endogenous murine tau protein isoforms and are not transgenic for any human tau. Infusion of AD_be_ increased Aβ load at the inoculation site as well as tau lesions that spread in connected areas notably the perirhinal/entorhinal cortex. AD_be_– inoculated animals displayed memory and synaptic impairments. Synaptic defects were correlated with neuritic tau lesions but not with other tau lesions or Aβ pathology. Neuroinflammation was evaluated as a complementary measure. Unexpectidly, synaptic density and microglial load were correlated.

## Methods

### Human brain collection and characterization

Frozen brain samples (parietal cortex) from eight sporadic AD patients as well as two age-matched control individuals were collected from a brain donation program of the NeuroCEB and the CNR-prion brain banks (Supplementary Table 1). Consent forms were previously signed by either the patients themselves or their next of kin, in accordance with French bioethics laws. AD patients had a classical evolving form of the pathology, characterized by a disease duration of 5 to 8 years (n=4) or a more rapidly evolving form characterized by a disease duration of 6 months to 3 years (n=4). No case of hippocampal sclerosis was reported and all brain samples were PrPSc negative. Samples from AD brains were also negative for α-synuclein and TDP-43. All brain tissues were assessed by immunohistochemistry, as previously described in Lam et al. 2021 [15]. Briefly, 4-μm-thick paraffin sections were cut, deparaffinized in xylene, successively rehydrated in ethanol (100, 90, and 70%) and rinsed under running tap water for 10 min before immunohistological staining. They were then incubated in 99% formic acid for 5 min, quenched for endogenous peroxidase with 3% hydrogen peroxide and 20% methanol, and washed in water. Sections were blocked at room temperature for 30 min in 4% bovine serum albumin (BSA) in 0.05 M tris-buffered saline, with 0.05% Tween 20, pH 8 (TBS-Tween, Sigma). They were then incubated overnight at +4 °C with the 6F3D anti-Aβ antibody (Dako, 1/200), polyclonal anti-tau antibody (Dako, 1/500), monoclonal anti-alpha-synuclein (LB509, Zymed, 1/250), polyclonal anti-TDP43 (Protein Tech Group, 1/1000) routinely used for β-amyloid, tau, alpha-synuclein and TDP43 detection, respectively. Sections were further incubated with a biotinylated secondary antibody for 25 min at room temperature, and the presence of the secondary antibody was revealed by a streptavidin–horseradish peroxidase conjugate using diaminobenzidine (Dako, Glostrup, Denmark). Sliced were counterstained with Harris hematoxylin.

Brain samples were also evaluated by biochemistry. For tau protein extraction, brain homogenates were sonicated on ice for 5 min, centrifuged for 5 min at 3,000 x g at +4 °C, diluted in 20 mM Tris/2% SDS and sonicated on ice for 5 min. For Aβ, Iba1 and GFAP protein extractions, brain homogenates were sonicated (6 strokes, cycle 0.5, 30% amplitude) in a lysis buffer at a final concentration of 50mM Tris-HCl pH 7.4, 150 mM NaCl, 1% Triton-X-100 supplemented with 1X protease inhibitors (Complete^TM^ Mini, EDTA-free Protease Inhibitor Cocktail, Roche) and 1/100 diluted phosphatase inhibitors (Phosphatase Inhibitor Cocktail 2, Sigma-Aldrich). Samples were centrifuged at 20,000 x g for 20 minutes at +4°C and the supernatant was collected for further use. Extracted samples were stored at -80°C after evaluation of total protein concentration by a BCA assay (Pierce^TM^). For tau characterization, samples were diluted to 1 μg/μL, diluted in 2X lithium dodecyl sulfate (LDS, Thermo Fisher Scientific) buffer with reducers and heated at +100 °C for 10 min. 15 μg of samples were loaded on a 12% Bis-TrisCriterion^TM^ gel (Bio-Rad) and migrated in MOPS buffer for 1 hour at 165 V on ice. After protein transfer on nitrocellulose sheets, migration and quality of the transfer were checked with a ponceau S staining. The membrane was saturated for 1 hour at room temperature, and was then incubated with the AT100 (pT212-pS214, Life technologies MN1060), 2H9 (pS422, 4BioDx 4BDX-1501), tau-Nter (12-21, LB lab-made) or tau-Cter (clone 9F6, LB lab-made) antibodies overnight at + 4 °C. A peroxidase coupled secondary anti-rabbit or anti-mouse antibody was then applied for 45 min at room temperature. Immunoblotting was revealed by ECL. GAPDH (Sigma 9545) was used as a loading control. For Iba1 and GFAP evaluations, extracted samples were denatured at +90°C for 5 min in a buffer containing 1X LDS (NuPAGE® LDS sample buffer, Invitrogen) and DTT 1X (NuPAGE® sample reducing agent, Invitrogen). 10 µg of denatured protein were loaded per well. Samples and molecular weight marker (Bio-Rad Precision Plus Protein^TM^ Dual Color standards) were loaded on 4-20% Criterion^TM^ TGX^TM^ gels (Bio-Rad) and migration was performed in a 1X tris-glycine buffer (Bio-Rad) at 120V for 1 hour. Proteins were then transferred to a nitrocellulose membrane using the Trans-Blot® Turbo^TM^ (Biorad) system. Migration and quality of the transfer were checked with a ponceau S staining. The membrane was then blocked with a TBS/0.1%Tween, 5% milk solution for 1 hour at room temperature, and incubated with the primary antibody Iba1 (Wako 1919741, 1/2000), GFAP (Dako Z0334, 1/5000) or actin (Sigma A2066, 1/5000) diluted in saturation buffer overnight at +4°C. After washing in TBS/0.1%Tween solution, the membrane was incubated with the appropriate secondary HRP-conjugate antibody diluted to 1/5000 in TBS/0.1%Tween for 1h at room temperature. The chemiluminescent signal was revealed using the Clarity western ECL (Bio-Rad) kit and the Chemidoc^TM^ MP (Bio-Rad) imaging system. Protein band intensities were quantified on the ImageJ software and normalized by the actin expression level. For amyloid-β protein quantification, all assay-specific material (pre-coated microtiter plate, buffers, antibodies, standard solutions) was provided in the V-PLEX kit Aβ Peptide Panel 1 (6E10) (MSD®). Human brain homogenates were diluted to 1/5 (Ctrl samples) or 1/10 (AD samples) in the dilution buffer. As described in the manufacturer’s protocol, the microtiter plate was blocked for 1 hour at room temperature with the appropriate buffer. After washing, 25µl of detection antibody and 25µl of diluted sample or standard were added in duplicate to the wells and incubated under continuous agitation for 2h at room temperature. Wells were washed and 150µl of reading buffer was added. Plate reading was performed with the MSD Sector Imager 2400 (model 1200) multiplex assay system. Aβ_1-38_, Aβ_1-40_ and Aβ_1-42_ quantifications were performed with the Discovery Workbench 4.0 MSD® software. Tau protein quantifications (total tau and phosphot-tau181) were performed according to the manufacturer’s protocol. Briefly, brain homogenates were diluted to 1/100 and 1/200 in the provided dilution buffer. 50µl of standards or samples, as well as 50µl of detection antibody solution were added to wells and incubated for 14 hours at +4°C. After washing, 100µl of 1X anti-rabbit IgG HRP solution was added for a 30 min incubation period at room temperature. 100µl of stabilized chromogen were then added to each well for 30 min at room temperature, in the dark. The reaction was stopped by adding 100µl of Stop solution and the plate was read at 450 nm within the hour. Data were analyzed with GraphPad Prism 7 using the 4PL method. All samples were tested in duplicates.

### Brain extracts preparation

Parietal cortex samples from each patient were individually homogenized at 10% weight/volume (w/v) in a sterile 1X Dulbecco’s phosphate buffer solution in CK14 soft tissue homogenizing tubes at 5000 rpm for 20 sec (Precellys®, Bertin technologies). They were then sonicated on ice for 5 sec at 40% amplitude and centrifuged at 3000g for 5 min at +4°C. The resulting supernatant was aliquoted in sterile polypropylene tubes and stored at − 80 °C until use. Ten percent individual brain extracts were thawed on ice and combined together according to thee groups: Control (n=2), AD patients with a disease duration of 5 to 8 years (n=4, AD1), or a disease duration of 6 months to 3 years (n=4, AD2). Aβ levels, total tau and phospho-tau181 as well as Iba1 and GFAP protein levels were assessed by biochemistry in each brain extract.

### Transgenic mice

Mouse experiments used the APP_swe_/PS1_dE9_ mouse model of amyloidosis (C57Bl6 background) [9]. Aβ plaques can be detected as early as 4 months of age in these mice. They increase in number and total area with age [9]. This model expresses endogenous murine tau protein isoforms and is not transgenic for any human tau. At the time of the inoculation of AD or Ctrl brain extracts in their brains, at 2 months of age, these mice did not have Aβ plaques. Animals were studied four or eight months after intracerebral inoculation of the brain extracts (at 4 and 8 months post-inoculation (mpi) respectively, *n_Ctrl_*=11 and 15, *n_AD1_*=14 and 15, *n_AD2_*=12 and 20). Wild-type littermates injected with the Ctrl brain sample were used as controls for the behavioral tests (at 4 and 8 mpi respectively, *n_WT_*=6 and 12). A cohort of 21 animals was studied by immunohistochemistry one month post-inoculation (*n_Ctrl_*=6, *n_AD1_*=7, *n_AD2_*=8). As the Aβ plaques and tau loads were similar between AD1 and AD2 inoculated animals at a given time point (see results), measures from AD1 and AD2 animals were pooled within a single AD_be_- inoculated group. All APP_swe_/PS1_dE9_ mice were born and bred in our center (Commissariat à l’Energie Atomique, centre de Fontenay-aux-Roses; European Institutions Agreement #B92-032-02). All animals were randomly assigned to the experimental groups using a simple procedure: they were identified using increasing numbers based on their birthdate. Animals with increasing numbers were alternatively assigned to the Ctrl (animal 1, 4, 7…), AD1 (animal 2, 5, 8…) and AD2 groups (animal 3, 6, 9…). Males were exclusively used in this study in order to optimize group homogeneity (as Aβ plaque load varies between males and females). Mice were injected during different inoculation sessions and each group was randomly inoculated at each session to avoid an “order of treatment” confounding effect. All experimental procedures were conducted in accordance with the European Community Council Directive 2010/63/UE and approved by local ethics committees (CEtEA-CEA DSV IdF N°44, France) and the French Ministry of Education and Research (A17_083 authorization given after depositing the research protocol and associated ethic issues), and in compliance with the 3R guidelines. Animal care was supervised by a dedicated in-house veterinarian and animal technicians. Humane endpoints concerning untreatable continuous suffering signs and prostrations were taken into account and not reached during the study. Animals were housed under standard environmental conditions (12-h light-dark cycle, temperature: 22 ± 1°C and humidity: 50%) with *ad libitum* access to food and water. The design and reporting of animal experiments were based on the ARRIVE reporting guidelines [8]. Sample size choice was based on previous experiments for Aβ induction in APP_swe_/PS1_dE9_ mice after inoculation of human brain (n = 8 at 4 mpi estimated with significance level of 5%, a power of 80%, and a two-sided test) [10] and increased to take into account uncertainties for new markers (tau lesion load, memory and synaptic changes). No animals were excluded from the study. SL was aware of initial group allocation, but further analyses (memory evaluations and post-mortem studies) were performed blinded.

### Stereotaxic surgery

Ten percent Ctrl, AD1 and AD2 brain extracts were thawed on ice. Extracts were then pooled together according to their group and the three resulting combined samples (Ctrl_be_, AD1_be_, AD2_be_) were sonicated (70% amplitude, 10 sec on/off; Branson SFX 150 cell disruptor sonicator, 3.17mm microtip probe Emerson, Bron) on ice in a sterile environment, extemporaneously before stereotaxic injection.

Two month-old APP_swe_/PS1_dE9_ and wild-type littermates were anesthetized by an intraperitoneal injection of ketamine (1mg/10g; Imalgène 1000, Merial) and xylazine (0.1mg/10g; 2% Rompun, Bayer Healthcare). Local anesthesia was also performed by a subcutaneous injection of lidocaine at the incision site (1 µl/g; 0.5% Xylovet, Ceva). Mice were placed in the stereotaxic frame (Phymep) and bilateral injections of brain samples were performed in the dorsal hippocampus (AP -2 mm, DV -2 mm, L +/- 1 mm from bregma). Using 34-gauge needles and Hamilton syringes, 2µl/site of sample were administered at a 0.2µl/min rate. After the injection, needles were kept in place for 5 more minutes before removal and the incision was sutured. The surgical area was cleaned before and after the procedure using povidone iodine (Vétédine, Vétoquinol). Respiration rate was monitored and body temperature was maintained at 37±0.5°C with a heating pad during the surgery. Anesthesia was reversed with a subcutaneous injection of atipamezole (0.25 mg/kg; Antisedan, Vetoquinol). Mice were placed in a ventilated heating box (25°C) and monitored until full recovery from anesthesia. Postoperative anticipatory pain management consisted in paracetamol administration in drinking water (1.45 ml/20ml of water; Doliprane, Sanofi) during 48 hours.

### Behavioral evaluations

A novel object recognition task in a V-maze was used to investigate cognition at 4 mpi or 8 mpi on brain-extract inoculated APP_swe_/PS1_dE9_ mice. Wild-type littermates injected with the Ctrl_be_ were used as controls for the tests. Mice were handled for 2 minutes per day, for 5 days prior to any test to prevent stress effects during tasks. Prior to each test, mice were habituated to the experimental room for 30 minutes. The experimenter was blind to mouse groups. Performances were recorded using a tracking software (EthoVision XT14, Noldus).

The V-maze arena consisted in two 6 cm-wide, 33.5 cm-long and 15 cm-high black arms forming a V shape and exposed to a 50 lux-lighting. The test was divided into three phases, each one separated by 24 hours. At the beginning of each session, mice were placed at the center of the arena, *i.e.* at the intersection of the arms. During the habituation phase (day 1), mice were free to explore the empty arena for 9 minutes. The distance travelled was automatically recorded as an indicator of their exploratory activity. For the training phase (day 2), two identical objects (bicolor plastic balls) were placed at the end of each arm. Exploratory activity was evaluated as the time spent exploring the objects (manually recorded) and the distance travelled during the 9-minute trial. On the test day (day 3), one familiar object (a bicolor plastic ball) was replaced by a novel one of a different shape and material (a transparent glass flask). Recognition was assessed using a discrimination index, calculated as follows:

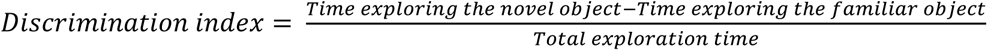

It reflects the time spent exploring each object, and therefore, the ability to discriminate a novel object from a familiar, previously explored one. A low discrimination index score reveals that mice spent less time exploring the new object, *i.e.* still had marked interest in the familiar object, and suggests that memory was impaired. Between each run, the V-maze was cleaned with 10% ethanol and dried.

### Animal euthanasia and brain preparation for histology

Mice were sacrificed at 4 or 8 mpi, after the behavioural tests, with an intraperitoneal injection of a lethal dose of pentobarbital (100 mg/kg; Exagon, Axience). They were perfused intracardiacally with cold sterile 0.1M PBS for 4 minutes, at a rate of 8 ml/min. The brain was extracted and post-fixed in 4% paraformaldehyde for 48 hours at +4°C, transferred in a 15% sucrose solution for 24 hours and in a 30% sucrose solution for 48 hours at +4°C for cryoprotection. Serial coronal sections of 40 µm were performed with a microtome (SM2400, Leica Microsystem) and stored at -20°C in a storing solution (glycerol 30%, ethylene glycol 30%, distilled water 30%, phosphate buffer 10%). Free-floating sections were rinced in a 0.1M PBS solution (10% Sigma-Aldrich® phosphate buffer, 0.9% Sigma-Aldrich® NaCl, distilled water) before use. Washing and incubation steps were performed on a shaker at room temperature unless indicated otherwise.

### Immunohistochemistry for Aβ, tau, microglia and astroglia

Aβ deposits were evaluated using a 4G8 labelling. Tau was evaluated using labelling with AT8 directed against hyperphosphorylated tau and AT100 that binds to a conformational epitope including phosphorylated Thr212 and Ser214. Microglia were evaluated using Iba1 and CD68 antibodies. Astrocytes were stained with the GFAP antibody.

4G8 labelling was performed after pretreating brain sections with 70% formic acid (VWR®) for 20 min at room temperature. AT8 and AT100 labellings were performed after a pretreatment with EDTA 1X citrate (Diagnostic BioSystems®) for 30 min at 95°C. All tissues were then incubated in hydrogen peroxide H_2_O_2_ 30% (Sigma-Aldrich®) diluted 1/100 for 20 min to inhibit endogenous peroxidases. Blocking of non-specific antigenic sites was achieved over 1 hour using a 0.2% Triton X-100/0.1M PBS (Sigma-Aldrich®) (PBST) solution containing 4.5% normal goat serum or 5% bovine serum albumin. Sections were then incubated at +4°C with the 4G8 (Biolegend 800706, 1/500), Iba1 (Wako 1919741, 1/1000), CD68 (Serotec – Biorad MCA 1957, 1/800) or GFAP (Dako Z0334, 1/10000) antibody diluted in a 3%NGS/PBST solution for 48h, or with the AT8 (Thermo MN1020B, 1/500) or AT100 (Thermo MN1060, 1/500) antibody diluted in a 3%NGS/PBST solution for 96h. After rinsing, an incubation with the appropriate biotinylated secondary antibody diluted to 1/1000 in PBST was performed for 1h at room temperature, followed by a 1h incubation at room temperature with a 1:250 dilution of an avidin-biotin complex solution (ABC Vectastain kit, Vector Laboratories®). Revelation was performed using the DAB Peroxidase Substrate Kit (DAB SK4100 kit, Vector Laboratories®). Sections were mounted on Superfrost Plus slides (Thermo-Scientific®). For the AT8 and AT100 labellings, a cresyl violet counterstain was performed. All sections were then dehydrated in successive baths of ethanol at 50°, 70°, 96° and 100° and in xylene. Slides were mounted with the Eukitt® mounting medium (Chem-Lab®).

Stained sections were scanned using an Axio Scan.Z1 (Zeiss® - Z-stack images acquired at 20× (z-stacks with 16 planes, 1µm steps with extended depth of focus)). Each slice was extracted individually in the .czi format using the Zen 2.0 (Zeiss®) software. Image processing and analysis were performed with the ImageJ software. Macros were developed for each staining in order to attain a reproducible semi-automated quantification. Images were imported with a 50% reduction in resolution (0.44 µm/pixel), converted to the RGB format and compressed in .Tif format. For the 4G8, Iba1 and CD68 immunostainings, segmentation was performed through an automatic local thresholding using the Phansalkar method (radius = 15). Aβ load was evaluated after quantification of the 4G8-labelled particles between 8 and 2,000 µm², and normalization to the surface area of each ROI. Microglial status was evaluated as a percentage of Iba1-or CD68-positive surface area in each ROI. For the AT8 and AT100 stainings, the blue component of each image was extracted to remove the cresyl violet counter-staining from the analysis. AT8-positive or AT100-positive tau loads were assessed following an automatic local thresholding of the staining with the Phansalkar method followed by an evaluation of the percentage of AT8-positive or AT100-positive surface area in each ROI.

In general, tau lesions occur in the form of neuropil threads, neurofibrillary tangles (NFTs), and neuritic plaques *i.e.* tau aggregates within neurites surrounding Aβ deposits. In addition for the AT8 immunostaining, a quantification of neuritic plaques and NFTs was performed by manual counting. The AT8-positive area stained within neuritic plaques was evaluated by drawing circular regions of interest (with a constant area of 6µm²), and by quantifying the percentage of tau-positive regions within each ROI, using the same thresholding method as previously described. A semi-quantitative analysis of neuropil threads was also performed by assigning a severity score based on the intensity and extent of AT8-positive staining in each ROI. All quantifications were performed on adjacent slices between -0.34 mm and -4.36 mm from bregma. Ten adjacent slices were analyzed for the 4G8 staining, and 5 for Iba1, CD68, AT8, and AT100 stainings. All ROIs were manually segmented using the Paxinos and Franklin neuro-anatomical atlas of mouse brain [21].

### Gallyas silver staining

Free-floating sections were mounted on Superfrost Plus (Thermo-Scientific®) slides and dried overnight prior to Gallyas staining. Section were permeabilized by successive incubations in toluene (2x5min) followed by ethanol at 100°, 90° and 70° (2 min per solution). Slides were then incubated in a 0.25% potassium permanganate solution for 15 min, in 2% oxalic acid for 2 min then in a lanthanum nitrate solution (0.04g/l lanthanum nitrate, 0.2g/l sodium acetate, 10% H2O2 30%) for 1h to reduce non-specific background. Several rinses with distilled water were performed followed by an incubation in an alkaline silver iodide solution (3.5% AgNO3 1%, 40g/l NaOH, 100g/l KI) for 2 min. The reaction was neutralized with 0.5% glacial acetic acid baths (3x1min) and sections were incubated for 20 min in a developing solution (2g/l NH4NO3, 2g/l AgNO3, 10g/l tungstosilicilic acid, 0.76% formaldehyde 37%, 50g/l anhydrous Na2CO3). Several rinses with 0.5% acetic acid (3x1min) followed by an incubation in 1% gold chloride solution for 5min were then carried out. Sections were rinsed with distilled water and the staining was fixed with a 1% sodium thiosulfate solution. All sections were then rinsed with distilled water and dehydrated for 1 to 5 min in successive baths of ethanol at 50°, 70°, 96° and 100° and in xylene. Slides were mounted with the Eukitt® mounting medium (Chem-Lab®). All steps were performed at room temperature.

### Co-stainings of microglia and Aβ plaques

In order to evaluate microglial load surrounding Aβ plaques, the co-staining of microglia and Aβ plaques was performed. Free-floating sections were permeabilized in a 0.2% Triton X-100/0.1M PBS (Sigma-Aldrich®) solution for 3x10min. Slices were stained by MXO4 dye (Tocris #4920, 1/300) for 30 min at room temperature, and then washed in a 0.1M PBS solution. Sections were blocked in a 4.5%NGS/PBST solution for 1h at room temperature before being incubated with the Iba1 antibody (Wako 1919741, 1/1000). Twenty four hours later, sections were rinsed in 0.1M PBS and incubated for 1h at room temperature with the appropriate secondary antibody diluted to 1/1000 in PBST (anti-rabbit AlexaFluor 633). Sections were rinsed and mounted on Superfrost Plus (Thermo-Scientific®) slides with the Vectashield® mounting medium with a refractive index of 1.45. Images of stained sections were acquired using a Leica DMI6000 confocal optical microscope (TCS SPE) with a 40x oil-immersion objective (refractive index 1.518) and the Leica Las X software. A confocal zoom of 3 and a pinhole aperture fixed at 1 Airy were applied. Acquisition was performed in sequential mode with a sampling rate of 1024x1024 and a scanning speed of 700 Hz. Image resolution was 60 nm/pixel and the optical section was 0.896 µm. Twelve separate planes with a 0.1 µm step were acquired. The excitation wavelengths were 633 nm (for Iba1) or 350 nm (for Aβ). Image acquisition was performed on 2 slices located between -3.28 mm and -4.24 mm from the bregma, with 3 images per slice for the CA1 region and for the perirhinal/entorhinal cortex. 3D deconvolution of the images was performed using the AutoQuant X3 software. The deconvoluted 8-bit images were analyzed using the ImageJ software. Quantification of microglial load around plaques was based on a thresholding procedure applied across all images to segment microglial cells. MXO4-positive surfaces were dilated as circular regions of interest (with a diameter of 40 µm) were drawn around the Aβ plaque to define a dilated plaque area. Microglial staining within the dilated surface, *e.g.* within the plaque area, was included in the analysis.

### Evaluation of synaptic density

Synaptic density was evaluated in the hippocampus (CA1) and the perirhinal/entorhinal cortex of all inoculated mice using a double immunolabelling of presynaptic (Bassoon) and postsynaptic (Homer1) markers. Free-floating sections were permeabilized in a 0.5% Triton X-100/0.1M PBS (Sigma-Aldrich®) solution for 15min. Slices were incubated with Bassoon (Abcam Ab82958, 1/200) and Homer1 (Synaptic systems 160003, 1/400) antibodies diluted in 3% BSA/PBST solution for 24 hours at +4°C. Incubation with secondary antibodies coupled to a fluorochrome (Alexa Fluor) diluted in a 3% BSA/0.1M PBS solution was then performed for 1h at room temperature. Sections were rinsed and mounted on Superfrost Plus (Thermo-Scientific®) slides with the Vectashield® mounting medium with a refractive index of 1.45. Images of stained sections were acquired using a Leica DMI6000 confocal optical microscope (TCS SPE) with a 63x oil-immersion objective (refractive index 1.518) and the Leica Las X software. A confocal zoom of 3 and a pinhole aperture fixed at 1 Airy were applied. Acquisition was performed in sequential mode with a sampling rate of 1024 x 1024 and a scanning speed of 700 Hz. Image resolution was 60 nm/pixel and the optical section was 0.896 µm. 26 separate planes with a 0.2 µm step were acquired. The excitation wavelengths were 594 nm or 633 nm. Image acquisition in the CA1 region was performed on 4 adjacent slices located between -1.82 mm and -3.28 mm from the bregma, with 2 images per slice. For the perirhinal/entorhinal cortex, 3 adjacent slices located between -3.28 mm and -4.24 mm from the bregma were analyzed, with 2 images acquired per slice. 3D deconvolution of the images was performed using the AutoQuant X3 software. The deconvoluted 8-bit images were analyzed using the ImageJ software, as described in Gilles et al [11]. Briefly, automated 3D segmentation of the staining allowed to determine the volume occupied by Bassoon-positive or Homer-positive objects in the 3D space as well as the localization of the geometrical centroid or center of mass of the objects. Co-localization was determined by the detection of overlapping objects, and depended on the center-to-center distance and the percentage of co-localization volumes for each pair of objects.

### Statistical analysis

Statistical analysis was performed using the GraphPad Prism software 8. For the behavioral tasks analysis, Kruskal-Wallis tests with Dunn’s multiple comparisons were perfomed except when repeated measures were acquired, in which case, a two-way repeated measures ANOVA with the Geisser-Greenhouse correction and Dunnett’s multiple comparisons was carried out. For the post-mortem analysis, Mann-Whitney tests were performed in order to compare differences between AD_be_- and Ctrl_be_-inoculated mice. For correlation studies, Spearman correlation test was performed. The significance level was set at *p*<0.05. Data are shown on scattered dot plots with mean ± standard error of the mean (s.e.m).

### Data availability

The data that support the findings of this study are available from the corresponding author, upon request.

## Results

### Characterization and inoculation of human brain homogenates

We prepared two brain homogenates from sporadic AD patients, with each homogenate consisting of a combination of four brain extracts from patients with a classical evolving form of the pathology (AD1) or a more rapidly evolving form (AD2). A third homogenate, considered as a control, was prepared from the brains of two non-demented individuals (Ctrl). The characteristics of the selected subjects are presented in Supplementary Table 1 and Supplementary Fig. 1. The amount of Aβ, tau and neuroinflammatory proteins slightly differed between the brain homogenates, as the AD2_be_ displayed more total tau and phospho-tau181, but less Aβ38 and Aβ40 than the AD1_be_ (Supplementary Fig. 1g-l). Iba1 and GFAP levels were similar in the two AD homogenates (Supplementary Fig. 1m-o).

### ADbe-inoculated Aβ transgenic mice develop memory alteration

AD_be_ and Ctrl_be_ were inoculated bilaterally in the dorsal hippocampus (CA1) of 2-month-old APP_swe_/PS1_dE9_ mice. An additional group of Ctrl_be_-inoculated wild-type littermates was used as an Aβ plaque-free control. The mice were evaluated in an object-recognition task (V-maze test) at 8 mpi. Animals from each group showed similar exploratory activity (similar distance travelled throughout the three days of test (Fig. 1a); comparable interest in the two identical objects during the training phase of the task (Fig. 1b)). During the memory task, *i.e.* the novel object recognition evaluation, Ctrl_be_-inoculated Aβ plaque-free wild-type and Aβ plaque-bearing APP_swe_/PS1_dE9_ mice had similar performances (Fig. 1c). This suggests that Aβ plaques did not modulate memory in this task. AD_be_-inoculated APP_swe_/PS1_dE9_ animals mice spent less time exploring the novel object than Ctrl_be_-inoculated mice. This suggests that AD_be_ infusion induces memory impairment (Fig. 1c). Another cohort of mice was also evaluated at 4 mpi. No difference in memory performance was observed at this stage (Supplementary Fig. 2a-c).

**Figure 1.**
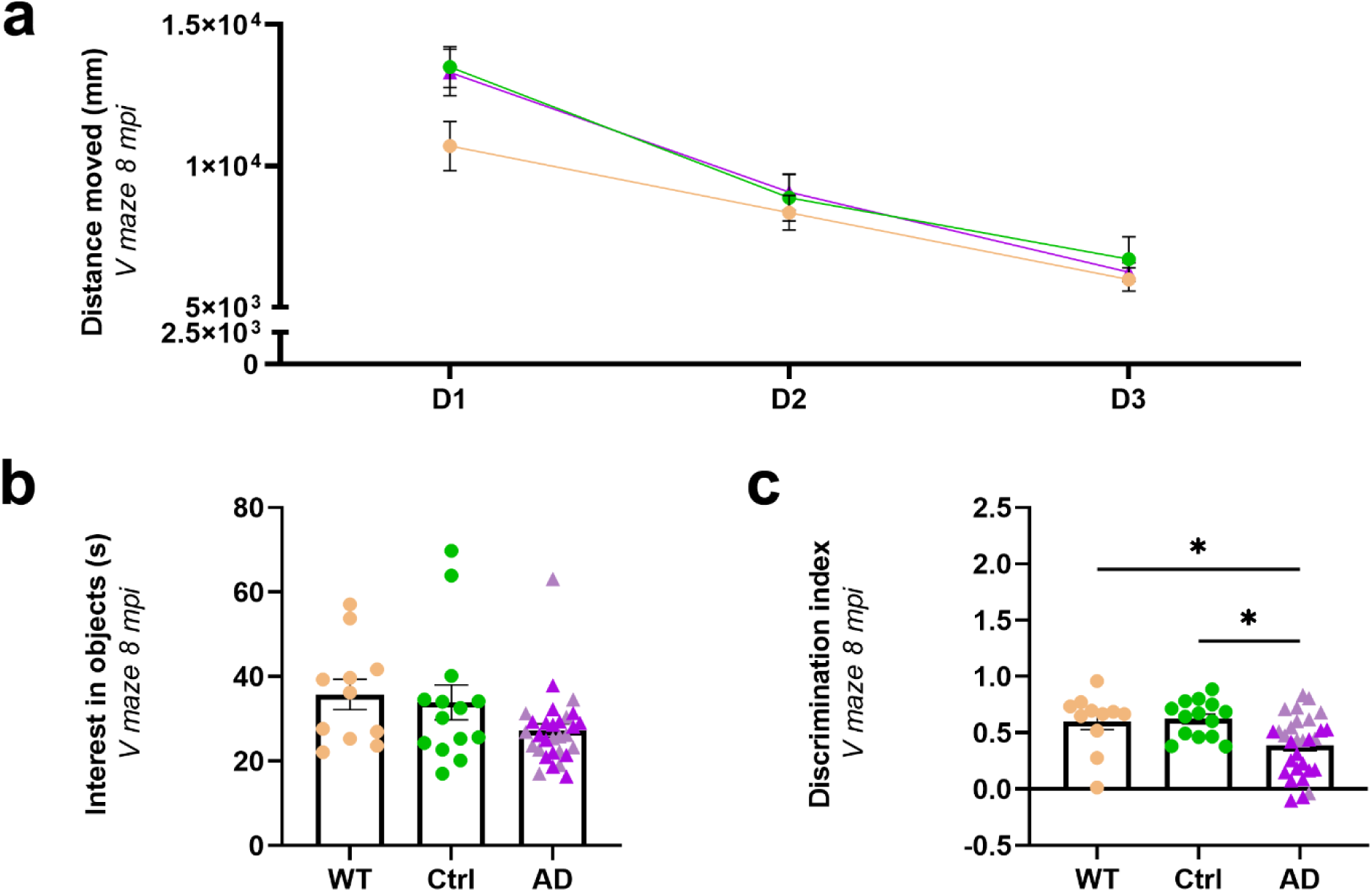
Object recognition deficits in the AD_be_-inoculated mice. Object recognition performances were evaluated at 8 mpi in a V-maze. WT mice and APP_swe_/PS1_dE9_ mice inoculated with Ctrl_be_ or AD_be_ had comparable exploratory activity, as suggested by the distance moved throughout the 3-day test (**a**, for the days: *F_(1.7, 90.5)_* = 72.4, *p*<0.0001; for the groups: *F_(2, 52)_* = 1.2, *p*=0.3; two-way repeated measures ANOVA with the Geisser-Greenhouse correction and Dunnett’s multiple comparisons) and the time spent on exploring the objects (**b**, *p*>0.05; Kruskal-Wallis with Dunn’s multiple comparisons). During the novel object recognition evaluation, AD_be_-inoculated mice spent less time exploring the novel object, as suggested by a lower discrimination index, compared to WT and APP_swe_/PS1_dE9_ Ctrl_be_-inoculated mice (**c**, respectively, *p*=0.027, and 0.016; Kruskal-Wallis with Dunn’s multiple comparisons). *n_Ctrl_*=15, *n_AD1_*=15 (light pink in b-c), *n_AD2_*=20 (dark pink in b-c), *n_WT_*=12 mice. \**p*<0.05; Data are shown as mean ± s.e.m.

### ADbe-infusion increases Aβ plaque deposition and tau pathologies close to the infusion site

At 8 mpi, infusion of AD_be_ led to an increased Aβ plaque deposition in the hippocampus and in the region surrounding the alveus compared to Ctrl_be_ (Fig. 2). The increase in Aβ pathology detected at 8 mpi already started at 4 mpi (Supplementary Fig. 2d-e).

**Figure 2.**
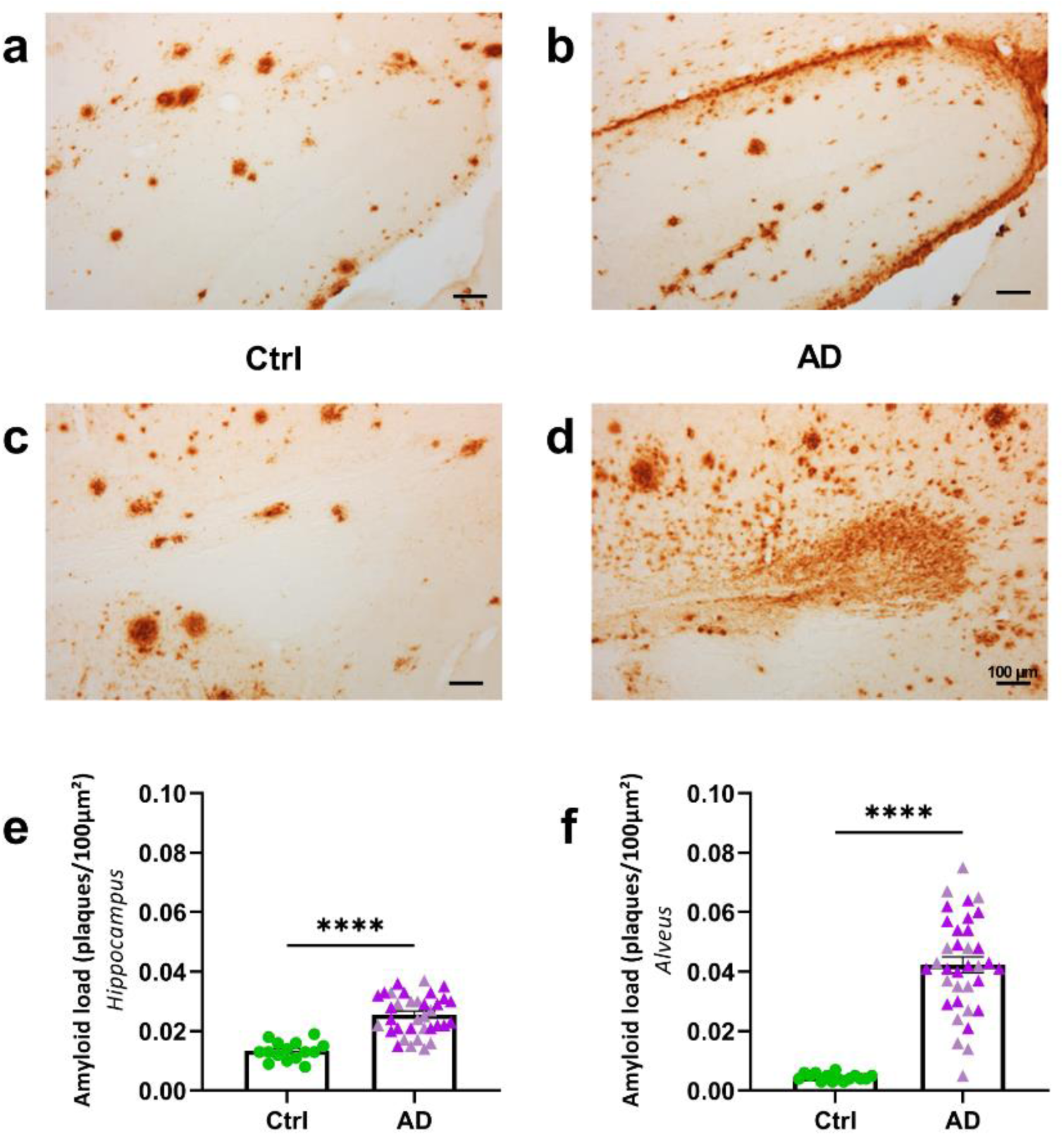
AD_be_ inoculation accelerates amyloidosis in APP_swe_/PS1_dE9_ mice close to the injection site. (**a-d**) Representative images of 4G8 immunolabelling showing Aβ pathology in the hippocampus (**a-b**) and alveus (**c-d**) of APP_swe_/PS1_dE9_ mice eight months after human brain inoculation. (**e-f**) Quantification of Aβ load (4G8-positive Aβ plaques per 100 µm²) revealed that AD_be_ inoculation accelerates Aβ deposition in the hippocampus (**e**, Mann Whitney’s test, U=21.5, *p*<0.0001) and alveus (**f**, Mann Whitney’s test, U=5, *p*<0.0001). \*\*\*\**p*<0.0001. *n_Ctrl_*=15, *n_AD1_*=15, *n_AD2_*=20 mice. Data are shown as mean ± s.e.m. Scale bars = 100 µm.

In humans, tau lesions occur in the form of neuropil threads, NFTs and neuritic plaques *i.e.* tau aggregates within neurites surrounding Aβ deposits. These three lesions were detected in the hippocampus of AD_be_-inoculated APP_swe_/PS1_dE9_ mice at 4 and 8 mpi (Fig. 3a: neuropil threads (NTs); Fig. 3b-d: NFTs; Fig. 3e: AT8-positive neuritic plaques (NPs)). At 1 mpi, tau lesions were not detected in any animal (Supplementary Fig. 3).

**Figure 3.**
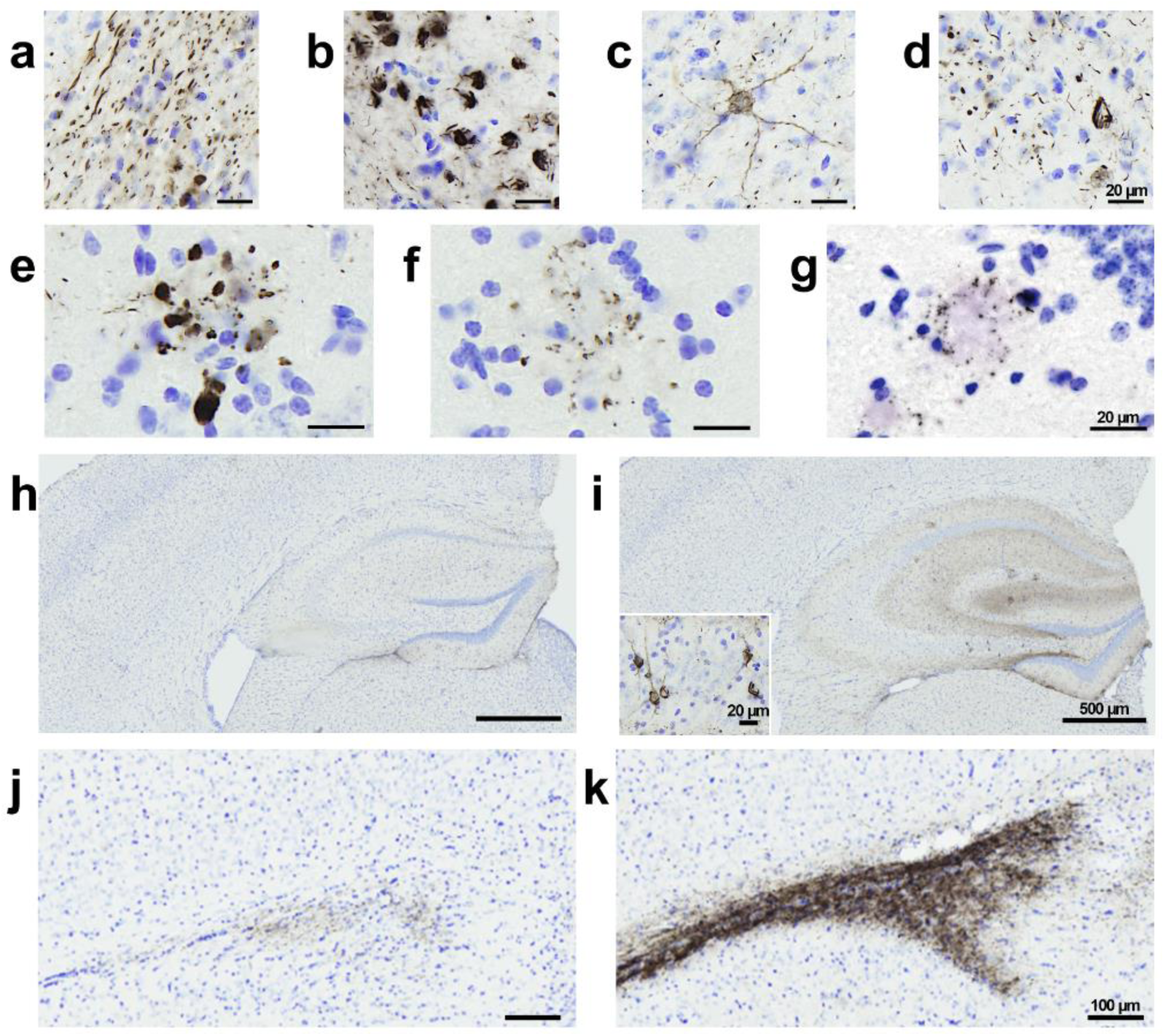
AD_be_ inoculation increases tau pathology in APP_swe_/PS1_dE9_ mice close to the injection site. AT8 immunolabelling revealed tau lesions in the forms of neuropil threads (**a**), NFTs (**b-d**), neuritic plaques (**e**) around the injection site of AD_be_-inoculated APP_swe_/PS1_dE9_ mice at 8 mpi. Neuritic plaques were also detected in Ctrl_be_-inoculated mice at 8 mpi (**f**). AT8 immunolabelling of non-inoculated aged APP_swe/_PS1_dE9_ mice (24 months old) also revealed neuritic plaques (**g**). (**h-i**) Representative images of AT8 immunolabelling showing tau pathology induction in the dorsal hippocampus of an AD_be_-inoculated APP_swe_/PS1_dE9_ mouse at 8 mpi (**i**) compared to a Ctrl_be_-inoculated littermate (**h**). (**j-k**) Representative images of AT8 immunolabelling showing tau pathology induction in the dorsal hippocampus of an AD_be_-inoculated APP_swe_/PS1_dE9_ mouse at 8 mpi (**k**) compared to a Ctrl_be_-inoculated littermate (**j**). Scale bars = 20 µm (**a-g**), 500 µm (**h-i**) and 100 µm (**j-j**) and 20 µm (**i insets**).

These three tau lesions did not occur similarly in AD_be_- and Ctrl_be_-inoculated APP_swe_/PS1_dE9_ mice. AT8-positive neuropil threads or NFTs were detected in the hippocampus of AD_be_- inoculated mice but not of Ctr_be_-inoculated animals (Fig. 4a-b). Neuropil threads were also detected in the alveus of AD_be_- but not of Ctr_be_-inoculated animals (Fig. 4c).

**Figure 4.**
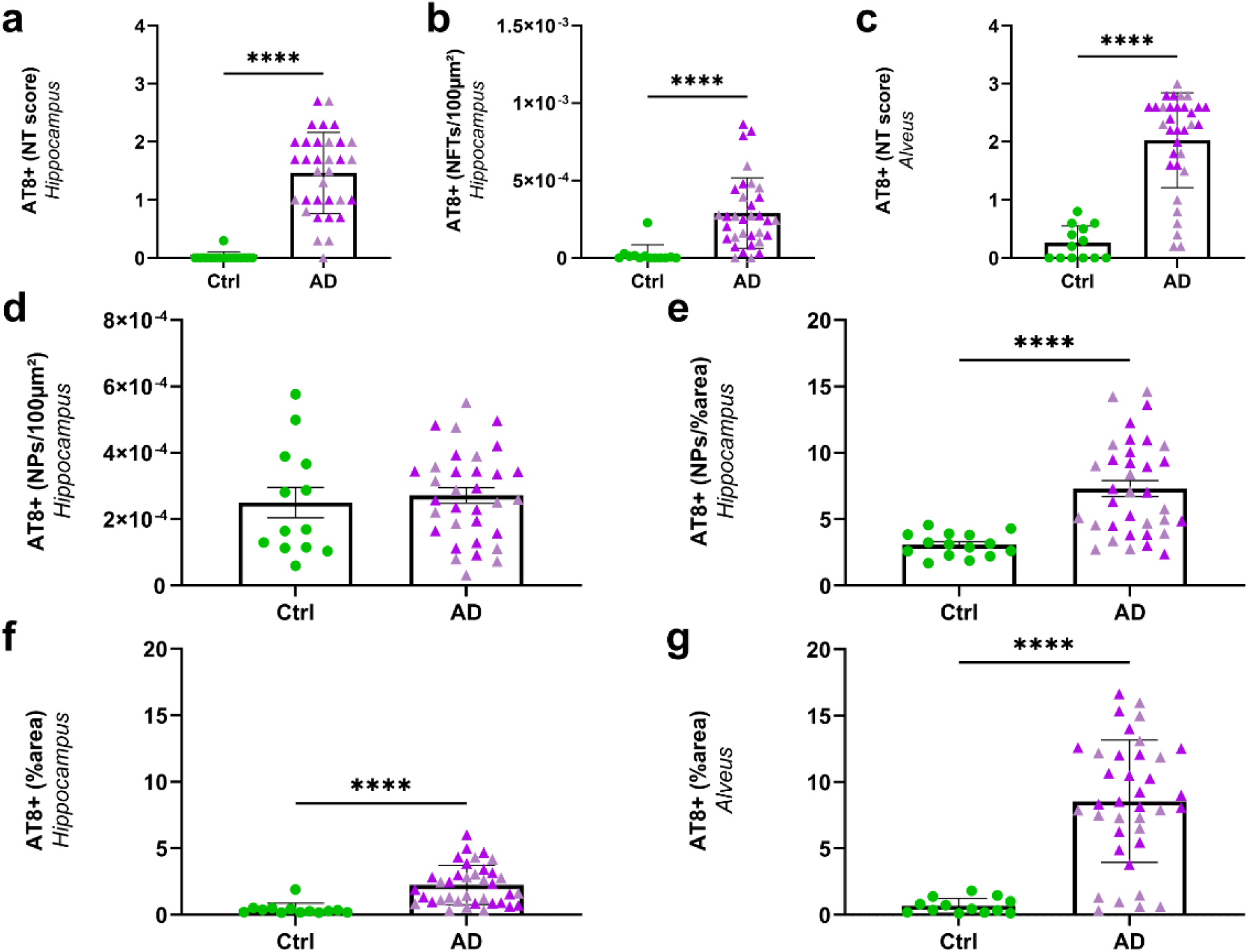
AD_be_ inoculation increases tau pathology in APP_swe_/PS1_dE9_ mice close to the injection site (8 mpi). Increased AT8-positive neuropil threads (**a**, *p*<0.0001) and NFTs (**b**, AT8-positive NFTs per 100 µm² of area, *p*<0.0001) in the hippocampus as well as neuropil threads in the alveus (**c**, *p*<0.0001) of AD_be_-inoculated animals compared to Ctrl_be_-animals. The number of AT8-positive neuritic plaque was similar in all groups (NPs per 100 µm² of area, **d**), but the AT8-positive area within neuritic plaques was higher in AD_be_- compared to Ctrl_be_- inoculated animals (**e**, *p*<0.0001). Increased overall AT8-labelled phospho-tau in the hippocampus (**f**) and alveus (**g**) following AD_be_ inoculation compared to Ctrl_be_ (percentage of AT8-positive area, *p*<0.0001). \*\*\*\**p*<0.0001. *n_Ctrl_*=15, *n_AD1_*=15, *n_AD2_*=20 mice. Mann Whitney’s tests. Data are shown as mean ± s.e.m. NP = neuritic plaques; NT = neuropil threads; NFT = neurofibrillary tangles.

Unlike neuropil threads or NFTs, neuritic plaques were detected in both AD_be_- (Fig. 3e) and Ctrl_be_-inoculated (Fig. 3f) APP_swe_/PS1_dE9_ mice at 8 mpi and the number of neuritic plaques was similar in their hippocampus (Fig. 4d). The AT8-positive area stained within neuritic plaques was however larger in the AD_be_-inoculated groups compared to Ctrl_be_ animals (Fig. 4e). To investigate whether human brain extract inoculation was necessary to induce neuritic plaques, we performed an AT8 immunostaining on old non-inoculated APP_swe_/PS1_dE9_ mice and observed neuritic plaques (Fig. 3g). Thus AT8-positive neuritic plaques occur spontaneously in APP_swe_/PS1_dE9_ mice.

Further quantitative analysis at 8 mpi showed that, compared to Ctrl_be_-inoculated mice, AD_be_- inoculated animals displayed an increase in AT8-positive tau lesions in the hippocampus (Fig. 3h-i, 4f) and in the alveus (Fig. 3j-k, 4g). This increase was already reported at 4 mpi, although to a lesser extent (Supplementary Fig. 2g-h).

AT100 detects late stage phospho-epitope found mostly in intracellular NFTs in humans and is used as a marker of aggregation [6]. It labelled neuropil threads and NFTs in AD_be_-inoculated mice (Supplementary Fig. 4a). AT100 staining was increased in the alveus of AD_be_-inoculated mice but not in their hippocampus (Supplementary Fig. 4b-c). Gallyas silver staining is used for the detection of aggregated tau pathology and paired helical filament-specific argentophilic tau lesions within neurofibrillary tangles. It revealed neuropil threads (Supplementary Fig. 4d), as well as Aβ plaques (Supplementary Fig. 4e) in AD_be_-inoculated mice.

### Spreading of tau pathology

We then evaluated Aβ and tau pathologies in the perirhinal/entorhinal cortex that is connected to the hippocampus (Fig. 5a-b). In this region, Aβ load was similar in AD_be_- and Ctrl_be_- inoculated APP_swe_/PS1_dE9_ mice at 8 mpi (Fig. 5c-e) and at 4 mpi (Supplementary Fig. 2f). Aβ detected in this region may thus only reflect the endogenous expression of the peptide in the APP_swe_/PS1_dE9_ model.

**Figure 5.**
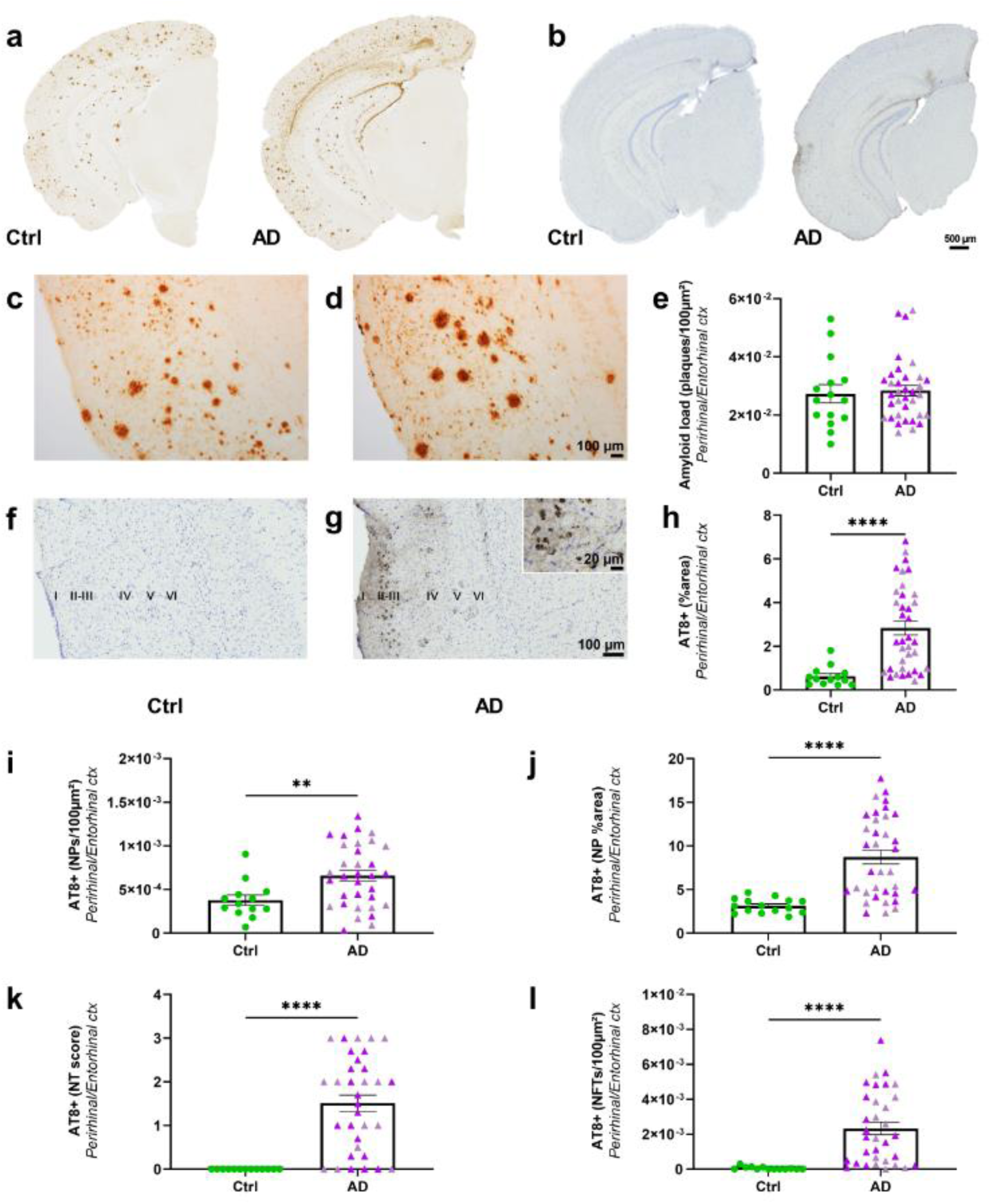
Aβ and tau pathologies in the perirhinal/entorhinal cortex following AD_be_ inoculation in APP_swe_/PS1_dE9_ mice (8 mpi). Representative images of the 4G8 immunolabelling showing Aβ pathology (**a**) and the AT8 immunolabelling showing tau pathology (**b**) in Ctrl_be_- and AD_be_-inoculated APP_swe_/PS1_dE9_ mice. Magnified views of Aβ (**c- d**) and tau (**f-g**) lesions in the perirhinal/entorhinal cortex. Quantification of Aβ load (**e**, *p*>0.05), overall AT8-positive tau lesions (**h**, *p*<0.0001), neuritic plaque (NPs) count (**i**, *p*=0.0098), AT8-positive area stained within neuritic plaques (**j**, *p*<0.0001), neuropil threads (NTs – **k**, *p*<0.0001), and NFTs (**l**, *p*<0.0001) in the perirhinal/entorhinal cortex. Mann Whitney’s tests. \*\**p*<0.01; \*\*\*\**p*<0.0001. *n_Ctrl_*=15, *n_AD1_*=15, *n_AD2_*=20 mice. Data are shown as mean ± s.e.m. Scale bars = 500 µm (**a-b**), 100 µm (**c-d, f-g**) and 20 µm in insert.

Tau pathology was detected in the perirhinal/entorhinal cortex, mainly in the external layers II and III of the cortex (Fig. 5f-g) that project to the dentate gyrus via the perforant pathway and to the CA1 region via the temporo-ammonic pathway, respectively. Internal layers (*e.g.* layers V-VI) that receive projections from the CA1 region were not labelled. At 8 mpi, areas of AT8- positive tau lesions were increased in the perirhinal/entorhinal cortex of AD_be_-inoculated mice compared to Ctrl_be_-inoculated ones (Fig. 5h). As for the hippocampus, the three categories of tau lesions did not occur similarly in AD_be_- and Ctrl_be_-inoculated APP_swe_/PS1_dE9_ mice. Indeed, neuritic plaques were detected in both AD_be_- and Ctrl_be_-inoculated mice. The number of neuritic plaques (Fig. 5i) and the AT8-positive area stained within these neuritic plaques (Fig. 5j) were however increased in AD_be_-inoculated mice compared to Ctrl_be_-inoculated ones. On the contrary, neuropil threads and NFTs only occurred in AD_be_ animals (Fig. 5k-l).

Other cortical regions such as the visual cortex also displayed neuropil threads, NFTs and neuritic plaques in AD_be_-inoculated mice (Supplementary Fig. 5a-b).

At 4 mpi, AT8-positive tau lesion loads were not different in the perirhinal/entorhinal cortex of AD_be_- and Ctrl_be_-inoculated mice (Supplementary Fig. 2i). The increased presence of tau in this region at 8 mpi in AD_be_-inoculated mice, thus reflects a time-dependent tau pathology spreading from 4 to 8 mpi.

AT100-positive tau lesions (Supplementary Fig. 6a-c) and Gallyas-positive (Supplementary Fig. 6d-e) tau lesions were detected in the perirhinal/entorhinal cortex of AD_be_-inoculated mice. Gallyas staining also revealed Aβ plaques (Supplementary Fig. 6d and 6f) in this region.

### Synaptic density is correlated with neuritic tau pathology but not with Aβ plaque load

We then evaluated synaptic density in the inoculated mice (Fig. 6). Synaptic density was decreased by 36% in the perirhinal/entorhinal cortex (Fig. 6c) and 19% in the CA1 of the hippocampus (Supplementary Fig. 7) of AD_be_-inoculated mice compared to Ctrl_be_ mice at 8 mpi. Synaptic density was not changed at 4 mpi (Supplementary Fig. 2j-k). We then focused on relationships between synaptic density and Aβ or tau lesions in the perirhinal/entorhinal cortex, *i.e.* at distance of the inoculation site at 8 mpi. Synaptic density was inversely correlated with neuritic plaque count (Fig. 6d) and AT8-positive area stained within neuritic plaques (Fig. 6e). Other tau-positive lesions (neuropil threads (Fig. 6f) or NFTs (Fig. 6g)) were not correlated with synaptic density. Synaptic density was neither correlated with Aβ load (Fig. 6h).

**Figure 6.**
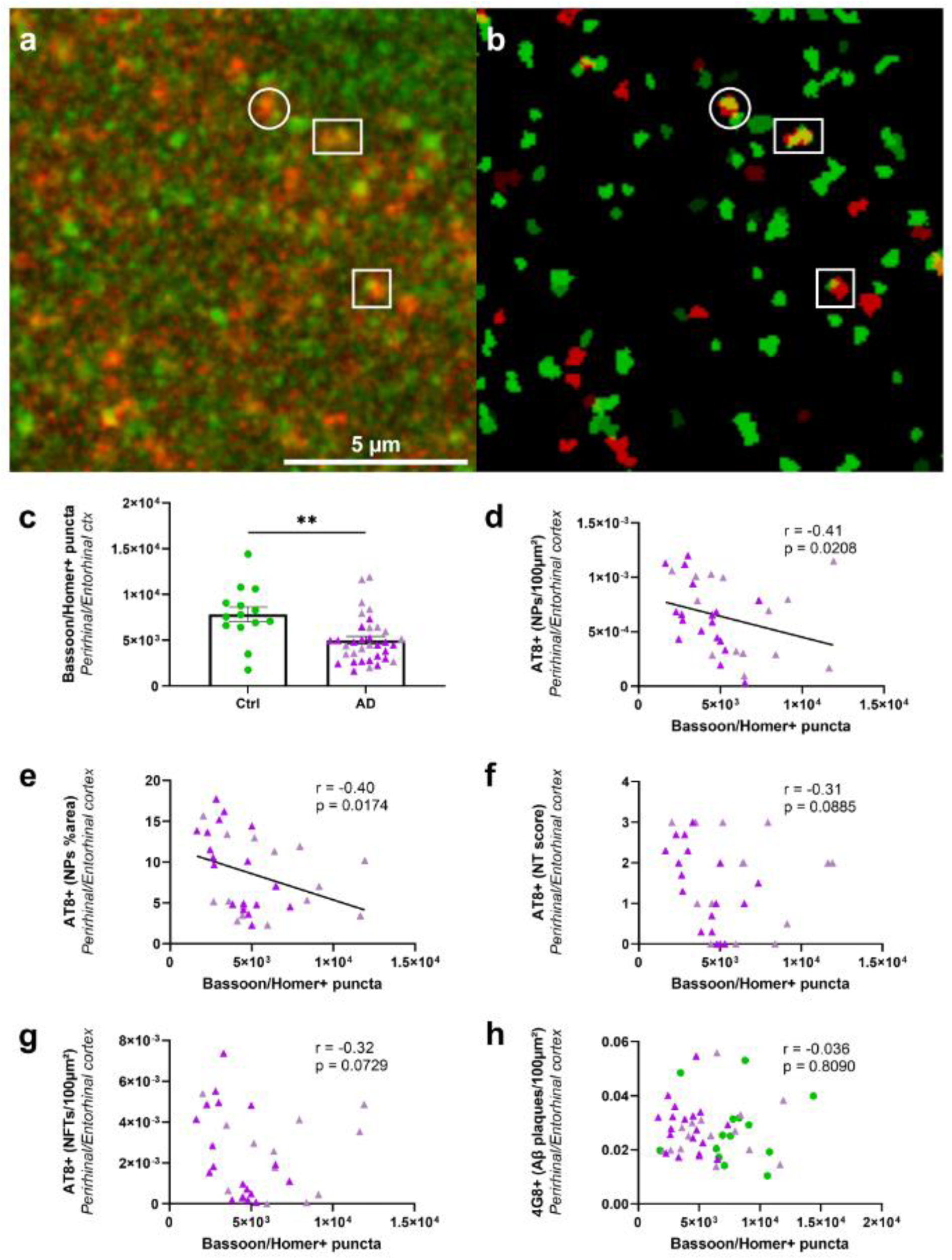
Synaptic density is decreased in AD_be_-inoculated APP_swe_/PS1_dE9_ mice and correlates with tau pathology in neuritic plaques (8 mpi). Representative image of Bassoon (red) and Homer-1 (green) immunolabelling (**a-b**). Bassoon/Homer-positive puncta (yellow in a-b) were quantified as an index of synaptic density. The synaptic density decreased in perirhinal/entorhinal cortex of AD_be_ mice compared to Ctrl_be_-inoculated mice at 8 mpi (**c**, *p*=0.002). Synaptic density was inversely correlated with tau pathology in neuritic plaques in the perirhinal/entorhinal cortex (**d**: neuritic plaque count; e: AT8-positive area within neuritic plaques), but not with neuropil threads (**f**), neurofibrillary tangles (**g**), or Aβ pathology (**h**). Spearman’s correlations with significance level at *p*<0.05. *n_Ctrl_*=15, *n_AD1_*=15, *n_AD2_*=20 mice. \*\**p*<0.01; Data are shown as mean ± s.e.m.

### Synaptic impairments are correlated with reduced microglial activation

Neuroinflammation was evaluated as a complementary endpoint. It was assessed by staining brain tissues using Iba1 antibody (Fig. 7a-f), a general marker for microglia as well as an anti- CD68 antibody (Fig. 7g-h) a marker of phagocytic microglia that stains a lysosomal protein expressed at high levels by activated microglia. Visual observation of the Iba1 stained sections suggested different levels of staining with some animals displaying high labelling (Fig. 7a-b) with abundant microglia with an activated phenotype characterized by beading with spheroidal swellings and enlarged cell body with dystrophic ramifications (Fig. 7c, arrows) [25]. Some other animals had lower staining (Fig. 7d-e) with highly ramified microglia (Fig. 7f).

**Figure 7.**
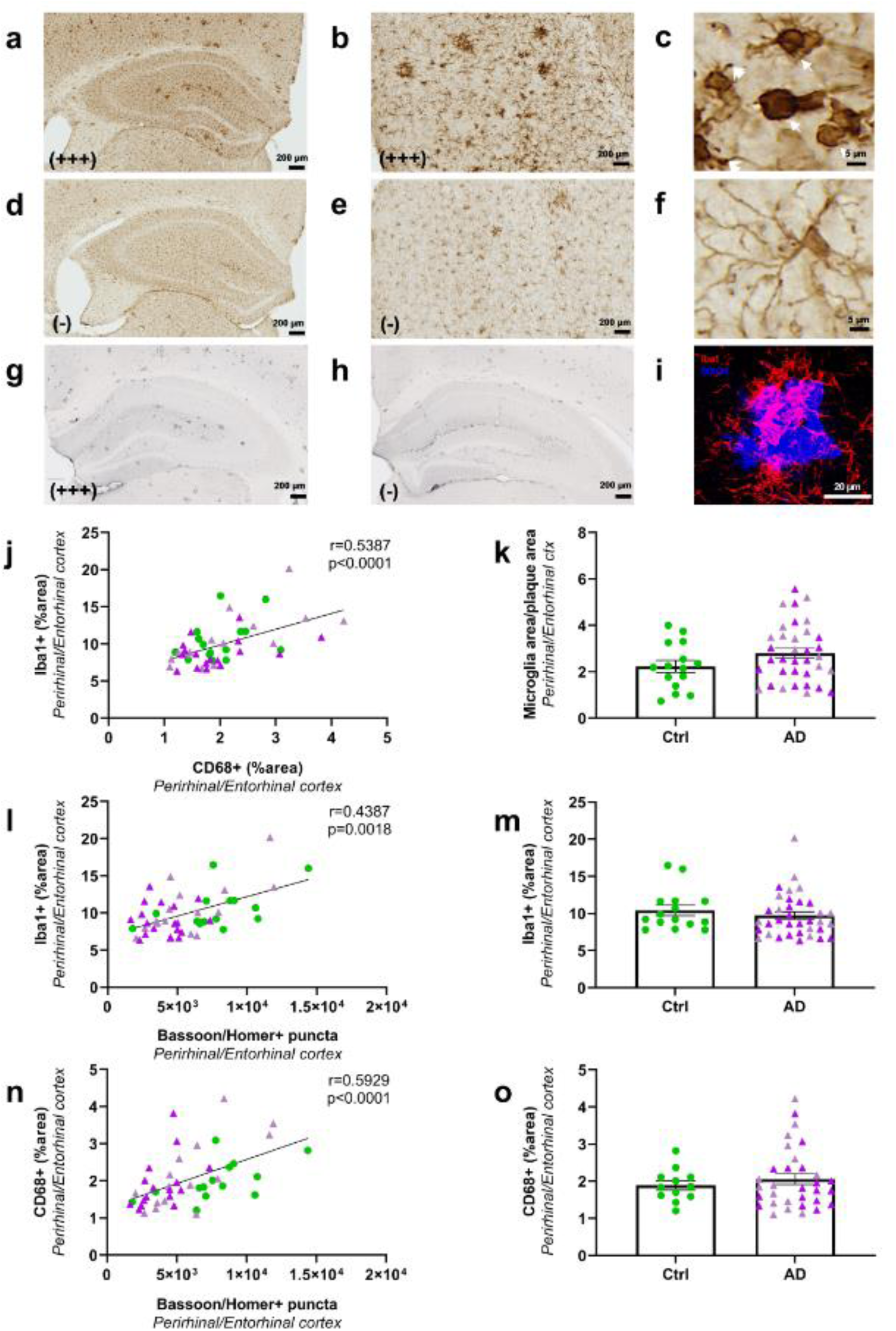
Microglial activation in AD_be_ and Ctrl_be_-inoculated APP_swe_/PS1_dE9_ mice (8 mpi). Within each group, animals with high and low Iba1 staining were identified. Representative images of one animal with higher Iba1 levels showing abundant staining in the hippocampus (**a**) and perirhinal/entorhinal cortex (**b**). Animals with higher Iba1 levels displayed activated microglia characterized by beading with spheroidal swellings of the processes and enlarged cell body with dystrophic ramifications (**c**, arrows). Animals with lower Iba1 levels (**d-e**) displayed more cells with a highly ramified profile (**f**) consistent with a non-activated phenotype. Representative images of one animal with higher (**g**) or lower (**h**) CD68 staining in the hippocampus. Iba1 staining level was correlated with CD68 labelling in the perirhinal/entorhinal cortex (**j**). Similar levels of Iba1 (**m**) and CD68 (**o**) stained areas in the perirhinal/entorhinal cortex of Ctrl_be_ and AD_be_ animals (Mann Whitney’s tests). Confocal microscopy showed Iba1-positive microglia (red) surrounding Aβ deposition stained by MXO4 (blue) (**i**). The Iba1-positive microglial load surrounding the Aβ plaques was similar in the different groups (**k**, Mann Whitney’s test). Interestingly, synaptic density was positively correlated with microglial Iba1 (**l**) or CD68 (**n**) labelling in the perirhinal/entorhinal cortex (Spearman’s correlations). *n_Ctrl_*=15, *n_AD1_*=15, *n_AD2_*=20 mice. Scale bar = 20µm.

Interindividual difference was also detected for CD68 labelling, with some animals displaying high labelling (Fig. 7g) and some others lower staining (Fig. 7h). Iba1 staining was positively correlated with CD68 staining in the hippocampus (r=0.39, p=0.009; not shown) or the perirhinal/entorhinal cortex (Fig. 7j).

AD_be_ and Ctrl_be_-inoculated mice had similar levels of Iba1 and CD68 stained areas in the hippocampus (not shown) or perirhinal/entorhinal cortex (Fig. 7m, 7o). Microglial is known to surround Aβ plaques where their activation shields Aβ plaques off from neurons to prevent their toxicity [5]. As expected, using confocal microscopy, we could detect Iba1-stained microglia surrounding Aβ plaques (Fig. 7i). We could not detect any differences in Iba1 staining around plaques in the different groups (Fig. 7k). This suggests no difference in the shielding effect in our inoculated groups. As for tau and Aβ, at 8 mpi, we further evaluated relationships between synaptic density and microglia in the perirhinal/entorhinal cortex, *i.e.* at distance of the inoculation site. A positive correlation was reported between synaptic density and Iba1 (Fig. 7l) or CD68 stainings (Fig. 7n).

Astrocyte reactivity (GFAP staining) was evaluated and no difference was detected between the groups in the hippocampus or in the perirhinal/entorhinal cortex (Supplementary Fig. 8). Also, synaptic density was not correlated with astrocyte-associated GFAP staining (not shown).

## Discussion

We infused AD_be_ into the brain of APP_swe_/PS1_dE9_ Aβ plaque-bearing mice. This induced tau lesions that spread in the brain, increased Aβ pathology close to the inoculation site, and induced downstream events including memory impairment and reduction of synaptic density. The time course of the induced pathology is reported at 1, 4 and 8 mpi (Supplementary Fig. 9). The three forms of tau lesions (neuritic plaques, neuropil threads and NFTs) were detected in AD_be_-inoculated animals. The lack of tau detection at one month post-inoculation and its progressive occurrence thereafter suggests that tau detected at 4 months and later does not reflect a direct deposit of human tau from AD_be_ and that it arises from mouse tau.

Neuritic plaques, but not the other tau-lesions, were detected in Ctrl_be_-inoculated animals as well as in aged non-inoculated APP_swe_/PS1_dE9_ mice. Thus Aβ aggregates, *per se*, are sufficient to induce tau lesions in surrounding neurites leading to neuritic plaques as reported in previous reports [17, 19, 23]. The density of tau lesions within neuritic plaques was increased at the inoculation site in AD_be_-inoculated animals, while the number of neuritic plaques was not increased at this level. This suggest that AD_be_ inoculation worsened the already present neuritic pathology.

Unlike neuritic plaques, neuropil threads and NFTs were only induced in AD_be_-inoculated animals. Two sources of tau seeds were available in AD_be_-inoculated animals: endogenous murine tau issued from neuritic plaques (detected in both Ctrl_be_- and AD_be_-inoculated animals), and exogenous tau seeds issued from human AD_be_. Because of the presence of neuropil threads and NFTs only in AD_be_-inoculated animals, we hypothesize that exogenous tau from AD_be_ are the main source of seeds for tau induction and spreading in neuropil threads and NFTs. Consistent with this hypothesis, in a previous study, we induced neuropil threads and NFTs in primates by infusing the AD_be_ used in the currend study [15]. We cannot totally rule out the hypothesis that endogenous tau seeds issued from neuritic plaques could participate to secondary tau pathology induction within neuropil threads and NFTs, but the presence of neuritic plaques in Ctrl_be_-inoculated animals that did not develop neuropil threads and NFTs, suggest that neuritic plaque are not sufficient for neuropil threads and NFTs induction in our model and that exogenous tau from AD_be_ are indeed the main source of seeds for neuropil threads and NFTs.

In the perirhinal/entorhinal cortex that is connected to the hippocampus, both the number of neuritic plaques and the AT8-positive area stained within these neuritic plaques were increased in AD_be_-inoculated animals. This suggests that increased tau lesions from the hippocampus participated to increased neuritic plaque occurrence and severity.

One can wonder whether amyloid pathology was required to induce tau pathology. Previous studies outlined that amyloid enhances the tau pathology induced by AD_be_-inoculation [31] but tau pathology can be induced in wild-type mice [1, 12], thus Aβ is not necessary for tau pathology induction.

Synaptic health has never been evaluated in models with Aβ and tau pathology induced by inoculation of human brain extracts. Tau pathology within neuritic plaque (plaque counts or AT8-positive area) in the perirhinal/entorhinal cortex, *i.e.* at distance from the inoculation site, was correlated with synaptic deficits while Aβ plaques or tau pathology within neuropil threads or NFTs were not corelated with synaptic changes. These results, in a model that is not based on tau overexpression, strongly supports the hypothesis that neuritic tau pathology contributes to synaptic impairment. Our results suggesting tau toxicity, however do not rule out a possible toxic role of Aβ on synaptic impairment. Indeed, AD_be_- and Ctrl_be_-inoculated mice from our study had similar Aβ plaque load in the perirhinal/entorhinal cortex, thus the Aβ plaque load effect can not be determined in our experimental set-up.

Memory evaluations did not show any difference between Ctrl_be_-inoculated Aβ plaque-free wild-type and Aβ plaque-bearing APP_swe_/PS1_de9_ mice at 8 mpi, *i.e.* at an age when Aβ plaques are numerous. This suggests a limited effect of Aβ plaques on memory at 8 mpi. On the contrary, AD_be_-inoculated animals that displayed tau lesions had memory impairments, which suggest a major impact of tau on the occurrence of cognitive changes. As Aβ and tau have been proposed to interact to induce memory changes [24], it is also possible that increasing tau pathology in mice increased Aβ impact on cognition.

Microglia was evaluated as a complementary measure. Although we did not detect differences of microglia load in AD_be_- and Ctrl_be_-inoculated animals, we decided to assess local relationships between microglia and synapses in the perirhinal/entorhinal cortex *i.e.* at distance from the inoculation site, in regions that did not receive the inoculates of brain extracts. Unexpectidly, we found a correlation between synaptic loss and a reduction of microgliosis (for both Iba1 and CD68 stainings). This is the first observation of a relationship between microglial activation and synapses density in a mouse model with Aβ and tau lesions without genetic or chemical manipulation of microglia. One possible interpretation is that the toxic process leading to synaptic loss also impaired microglia. Tau pathology can drive microglial degeneration [25, 29]. As we found a relationship between neuritic tau pathology and synaptic deficit, we can propose the hypothesis that tau could be the toxic species. This hypothesis needs now to be formally tested (for exemple with treatments that inhibit microglia as CSF1R inhibitors [20]). Finally, one can wonder what are the key compounds present in the human brain extracts to induce the tau and synaptic pathology: Aβ, soluble tau, fragments of fibrils from each, other factors such as cytokines, etc? Answering to this question is beyond the scope of this article. Several previous studies have shown that exogenous tau can transmit tau pathology [4], but this does not rule out the possible effects of other compounds on tau pathology induction. Regarding synaptic pathology, at the inoculation site, several compounds from the AD_be_ could participate to synaptic changes. By studying synaptic impairments in the perirhinal/entorhinal cortex, *i.e.* at distance from the inoculation site (the hippocampus) and in a region where tau was induced after spreading from hippocampal region, we expect that the effects of unspecified compounds from the AD_be_ was eroded. This reinforces the validity of the assumption that murine neuritic tau pathology participated to synaptic impairments, but still does not totally rule out the possible effects of other undetermined compounds.

## Conclusions

Intracerebral infusion of AD_be_- into Aβ plaque–bearing mice that do not overexpress tau induced memory and synaptic impairments, increases Aβ load at the inoculation site as well as tau lesions that spread in connected areas. Based on our results, we thus propose the following sequence of events. Aβ plaques can induce tau lesions within neuritic plaques. In the presence of exogenous tau seeds issued from human brain extracts, the tau pathology is amplified and leads to neuropil threads and NFTs. These events favor the cerebral spreading of tau pathology in the form of neuritic plaques, neuropil threads and NFTs. The neuric plaques, but not as much the other tau lesions, induce a reaction that is associated with synaptic density reduction. Microglia regulates or is regulated by events leading to synaptic reduction.

## List of abbreviations

Aβ: amyloid-β
AD: Alzheimer’s disease
AD_be_: AD brain extracts
Ctrl: control
Ctrl_be_: control brain extracts
mpi: months post-inoculation
NFTs: neurofibrillary tangles
NP: neuritic plaques
NT: neuropil threads
WT: wild-type

## Declarations

### Ethics approval and consent to participate

All experimental procedures were conducted in accordance with the European Community Council Directive 2010/63/UE and approved by local ethics committees (CEtEA-CEA DSV IdF N°44, France) and the French Ministry of Education and Research (A17_083 authorization given after depositing the research protocol and associated ethic issues), and in compliance with the 3R guidelines. Animal care was supervised by a dedicated veterinarian and animal technicians. Humane endpoints concerned untreatable continuous suffering signs and prostrations were used and not reached during the study.

### Consent for publication

Does not apply to the content of this article.

### Availability of data and materials

The data that support the findings of this study are available from the corresponding author, upon reasonable request.

### Author’s contributions

S.L., A.S.H., F.P., and M.D. contributed to the study conception and design. N.N.N., C.D. provided the human brain samples. N.N.N., S.L., S.B., C.D. and S.H. characterized the human brain samples. S.L., M.G. and M.G. performed the inoculations in mice. S.L. and K.C. designed and performed memory evaluations, A.S.H., F.P., and S.L. designed and performed the immunohistological analysis in animals. A.S.H., S.E., L.B., and S.L. performed biochemical analysis. S.L., A.S.H., and M.D. wrote the manuscript. All authors commented on previous versions of the manuscript. All authors read and approved the final manuscript.

## Acknowledgements

We thank Martine Guillermier and Mylène Gaudin for surgical expertise during inoculation of brain extracts to animals. We thank Nicolas Heck for his help in synapse quantification, and Cecilia Garrec for the editing and English language correction, and Nicolas Sergeant for a critical review of this article. We thank the donors and the Brain Donation Program of the NeuroCEB Brainbank Neuropathologist’s Network run by a consortium of Patient Associations: ARSLA (association for research on amyotrophic lateral sclerosis), CSC (cerebellar ataxias), Fondation ARSEP (association for research on multiple sclerosis), France DFT (fronto-temporal dementia), Fondation Vaincre Alzheimer, France Parkinson, with the support of Fondation Plan Alzheimer and IHU A-ICM for providing the brain samples used in this study. NeuroCEB Brainbank Neuropathologit’s Network members: Franck Letournel (Angers), Marie-Laure Martin-Négrier (Bordeaux), Maxime Faisant (Caen), Catherine Godfraind (Clermont-Ferrand), Jean Boutonnat (Grenoble), Claude-Alain Maurage (Lille), Vincent Deramecourt (Lille), Mathilde Duchesne (Limoges), David Meyronet (Lyon), Tanguy Fenouil (Lyon), André Mauès de Paula (Marseille), Valérie Rigau (Montpellier), Fanny Vandenbos-Burel (Nice), Danielle Seilhean (Paris), Charles Duyckaerts (Paris), Susana Boluda (Paris), Isabelle Plu (Paris), Dan Christian Chiforeanu (Rennes), Annie Laquerrière (Rouen), Florent Marguet (Rouen), Béatrice Lannes (Strasbourg), Benoît Lhermitte (Strasbourg).

## Supplementary data

**Supplementary Table 1.** Patient characteristics.

| Patient |  |  |  |  | Neuropathology |  |  |  |  |  |  |
| --- | --- | --- | --- | --- | --- | --- | --- | --- | --- | --- | --- |
| ID | Age | Gender | Disease progression (months) | Post mortem delay (hours) | Braak stage | Thal stage | CAA | α-synuclein | TDP43 | Hippocampal sclerosis | PrPSc |
| Ctrl1 | 66 | M | - | 26 | II | 0 | 0 | 0 | 0 | 0 | 0 |
| Ctrl2 | 76 | M | - | NA | II | 0 | 0 | 0 | 0 | 0 | 0 |
| AD1a | 79 | F | 78 | 48.6 | V | 5 | Type2 | 0 | 0 | 0 | 0 |
| AD1b | 87 | F | 72 | 29 | V-VI | 4 | Type1 | 0 | 0 | 0 | 0 |
| AD1c | 89 | F | 96 | 30.6 | V | 5 | Type1 | 0 | 0 | 0 | 0 |
| AD1d | 71 | M | 66 | 54 | VI | 4 | Type2 | Amygdala | 0 | 0 | 0 |
| AD2a | 84 | F | 6 | 79 | V | 4 | Type2 | 0 | 0 | 0 | 0 |
| AD2b | 81 | F | 36 | NA | V | 5 | Type1 | 0 | 0 | 0 | 0 |
| AD2c | 81 | F | 36 | 26 | VI | 4 | 0 | 0 | 0 | 0 | 0 |
| AD2d | 86 | F | 36 | 20.5 | V | 4 | Type2 | 0 | 0 | 0 | 0 |

Age-matched classical slowly (AD1) and rapidly evolving (AD2) Alzheimer patients were selected based on disease duration (over or under 36 months) and neuropathological evaluation, including similar Braak and Thal stages. Brains were negative for α-synuclein, TAR DNA-binding protein 43 (TDP43), hippocampal sclerosis and pathological prion PrPSc. Two non-Alzheimer control individuals (Ctrl) were also included in this study. NA: not available.

**Supplementary Figure 1:**
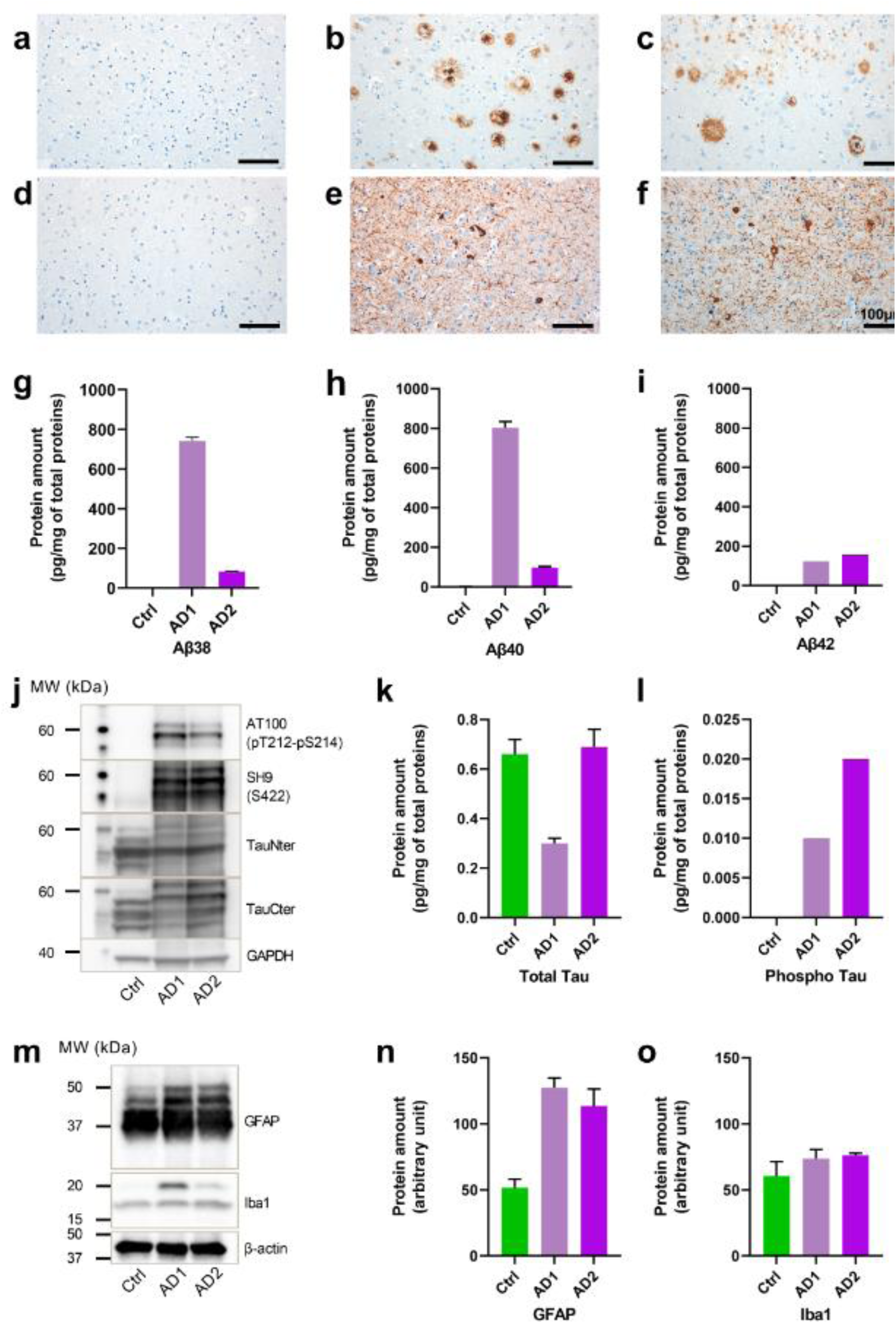
Characteristics of human brain samples and brain extracts inoculated to animals. Representative images of Ctrl, AD1 and AD2 brain samples stained for Aβ (a-c) and tau (d-f) pathologies. Scale bars = 100 µm. Three brain extracts were prepared from 2 control individuals, 4 cases with a slowly evolving form of AD and 4 cases with a rapidly evolving form of AD (Ctrl, AD1 and AD2 brain extracts, respectively). All quantifications of the brain extracts (Ctrl, AD1 or AD2) were performed in duplicate and data are shown as mean ± standard deviation of the replicates. (g-i) Quantifications of total Aβ38, Aβ40 and Aβ42 of the brain extracts (MSD technology). Both AD brain extracts had more Aβ proteins compared to the Ctrl one. The AD1 extract showed more Aβ38 and Aβ40 than the AD2 one. (j-l) Tau profile evaluation by western blot revealed a pathological hyperphosphorylated tau triplet at 60, 64 and 69 kDa observed in AD and a typical shift in the molecular weight of the Alzheimer Tau-Cter triplet in AD1 and AD2 brain extracts (j). Total tau (k) and pathological phospho-tau 181 levels (l) were assessed using ELISA quantification. Neuroinflammatory profile evaluation by western blots revealed higher astrocytic presence (GFAP-positive) in AD1 and AD2 brain extracts compared to the Ctrl extract (m-n). Microglial (Iba1-positive) levels were similar in the Ctrl, AD1 and AD2 groups (m, o). (data already published in [15]).

**Supplementary Figure 2.**
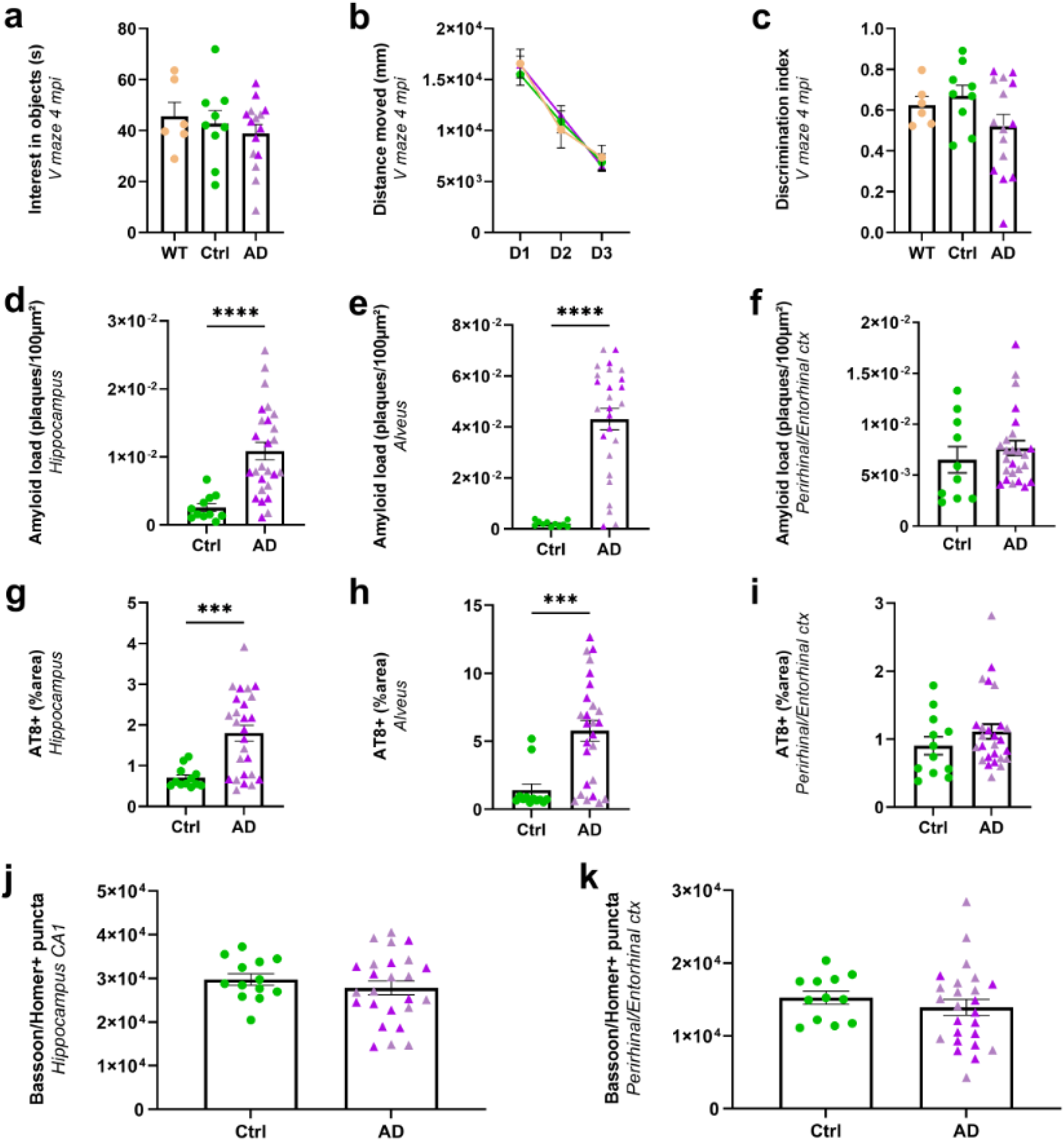
Cognitive performance, Aβ and tau loads evaluations, synaptic density at 4 mpi. (**a-c**) Object recognition performances were evaluated at 4 mpi using a V-maze test. WT mice and APP_swe_/PS1_dE9_ mice inoculated with Ctrl_be_ or AD_be_ had comparable exploratory activity, as suggested by the time spent on exploring the objects (**a**) (*p*>0.05; Kruskal-Wallis with Dunn’s multiple comparisons) and the distance moved throughout the 3-day test (**b**) (for the days: *F_(1.9, 51.7)_*=131.2, *p*<0.0001; for the groups: *F_(2, 27)_*=0.06, *p*=0.9; two-way repeated measures ANOVA with the Geisser-Greenhouse correction and Dunnett’s multiple comparisons). No difference in the novel object recognition test was reported between the groups, as similar discrimination indexes were observed (**c**) (*p*>0.05; Kruskal-Wallis with Dunn’s multiple comparisons). (**d-f**) Aβ load quantification at 4 mpi revealed that AD_be_ inoculation accelerates Aβ deposition in the hippocampus (**d**, *p*<0.0001; Mann Whitney’s test) and alveus (**e**, *p*<0.0001), but not the perirhinal/entorhinal cortex (**f**, *p*=0.3). (**g-i**) AT8-positive tau overall quantification at 4 mpi revealed that AD_be_ inoculation induces tau lesions in the hippocampus (**g,** *p*= 0.0007), in the alveus (**h**, *p*= 0.0003) but not the perirhinal/entorhinal cortex (**i**, *p*=0.2). (**j-k**) Quantification of Bassoon and Homer colocalization at 4 mpi did not show any differences in the CA1 (**j**) and in the perirhinal/entorhinal cortex (**k**) between the groups (*p*>0.05; Mann Whitney’s test). \*\*\**p*<0.001; \*\*\*\**p*<0.0001. *n_Ctrl_*=9, *n_AD1_*=7, *n_AD2_*=8, *n_WT_*=6 mice in a-c, *n_Ctrl_*=11, *n_AD1_*=14, *n_AD2_*=12 in d-k. Data are shown as mean ± s.e.m.

**Supplementary Figure 3.**
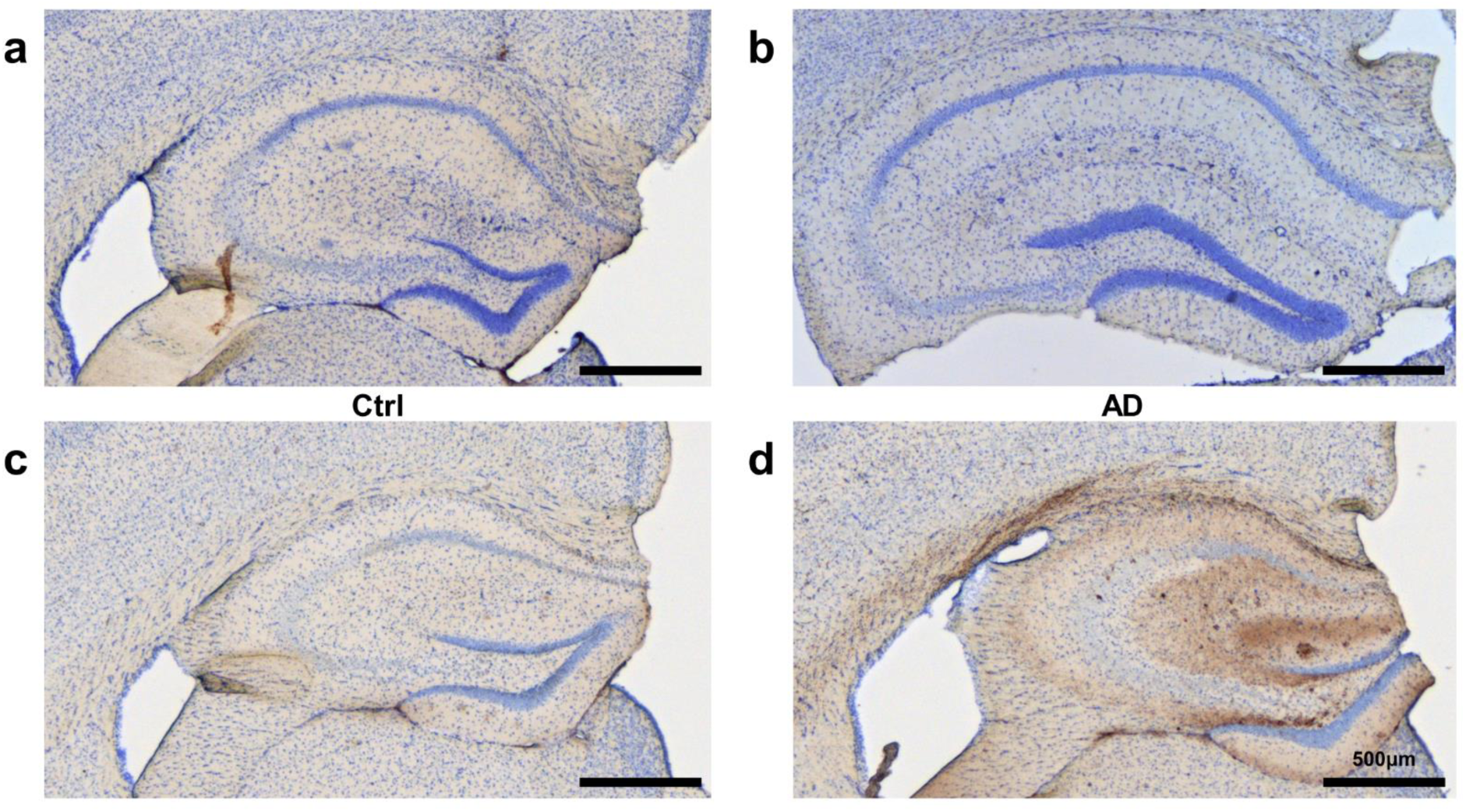
AT8 immunolabelling in the hippocampus of Ctrl_be_- (a, c) and AD_be_-inoculated (b, d) APP_swe_/PS1_dE9_ mice at 1 mpi (a-b) and 8 mpi (c-d). AT8-positive tau lesions were not detected in 1 mpi APP_swe_/PS1_dE9_ mice (b) while they were detected at 8mpi (d).

**Supplementary Figure 4.**
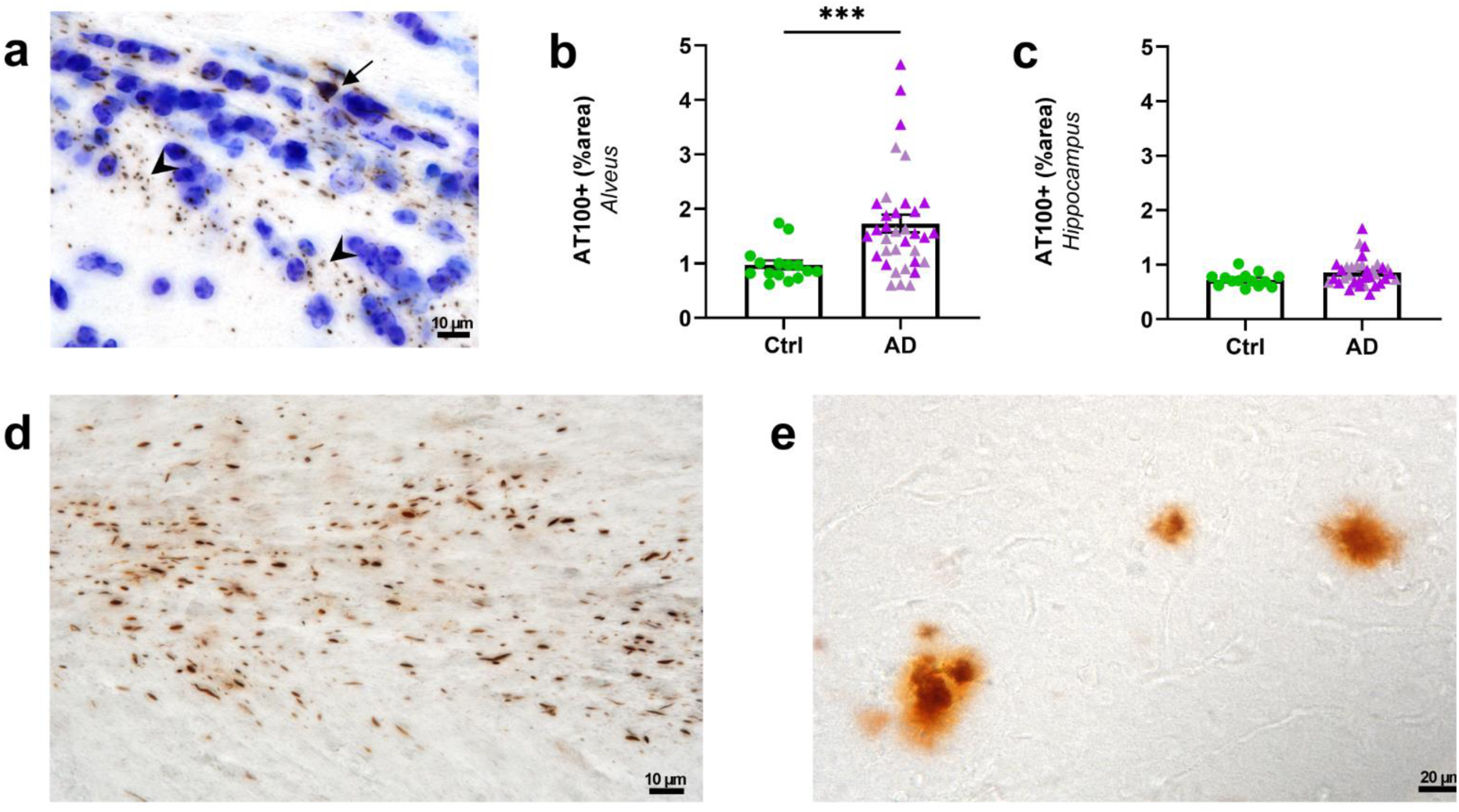
AT100 and Gallyas-positive lesions next to the inoculation site of APP_swe_/PS1_dE9_ mice 8 months after AD_be_ infusion. (**a**) AT100 staining revealed neuropil threads (arrowheads) and NFTs (arrow). (**b-c**) Overall quantification of AT100 labelling showed increased labelling in the alveus (**b**) and a trend for increased labelling in the hippocampus (**c**) of AD_be_-inoculated mice (*p*=0.0007 and 0.092 respectively, Mann-Whitney test). Gallyas silver staining revealed neuropil threads (**d**) as well as Aβ plaques (**e**) in AD_be_- inoculated mice. *n_Ctrl_*=15, *n_AD1_*=15, *n_AD2_*=20. Data are shown as mean ± s.e.m. Scale bars = 10 µm (**a, d**) and 20 µm (**e**).

**Supplementary Figure 5.**
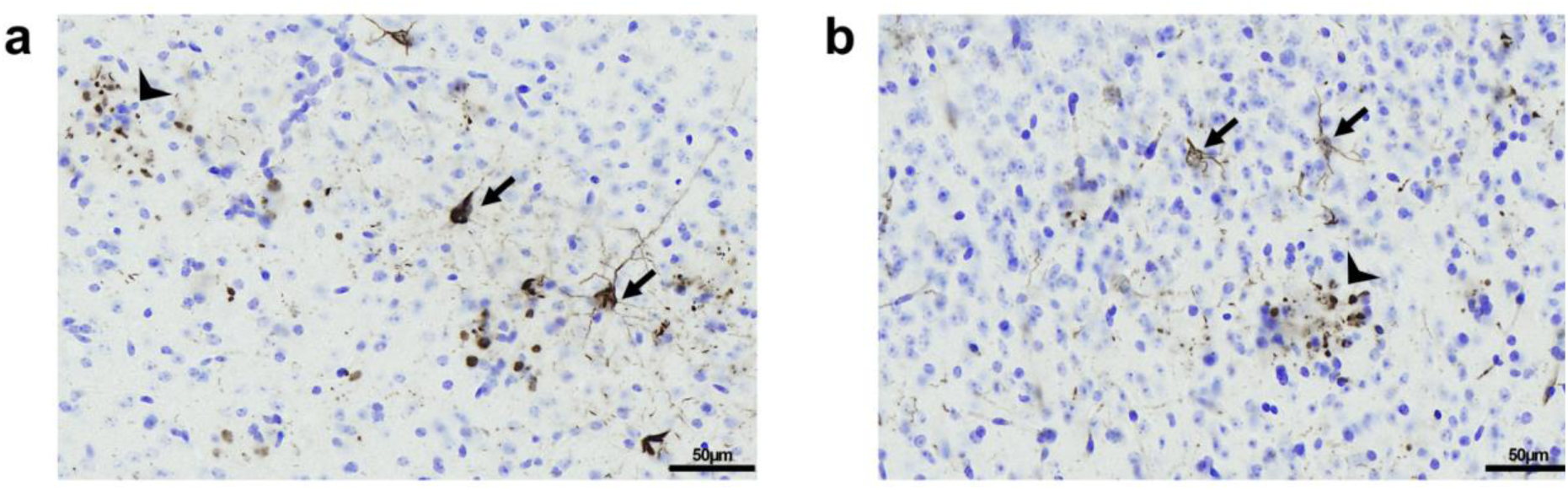
AT8-positive lesions in the visual cortex of APP_swe_/PS1_dE9_ mice 8 months after AD_be_ infusion. AT8 staining revealed NFTs (arrows) surrounded by neuropil threads, as well as neuritic plaques (arrowheads), in the visual cortex of AD brain-inoculated mice (**a**, **b**). Scale bars = 50 µm.

**Supplementary Figure 6.**
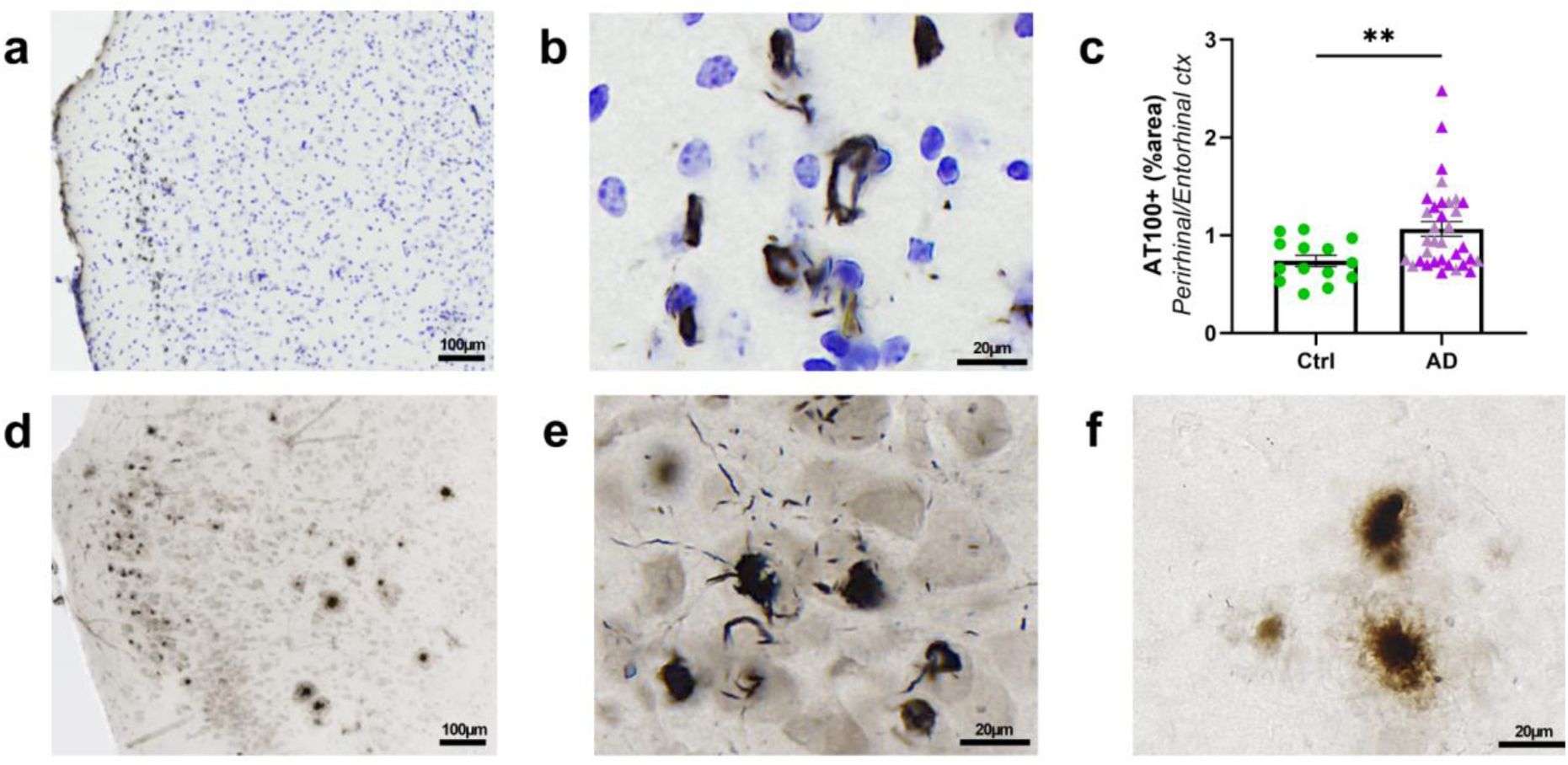
AT100- and Gallyas-positive lesions spreading to the perirhinal/entorhinal cortex of APP_swe_/PS1_dE9_ mice 8 months after AD_be_ infusion. (**a-b**) AT100 staining revealed NFTs in the perirhinal/entorhinal cortex. (**c**) Overall quantification of AT100 labelling showed increased labelling in the perirhinal/entorhinal cortex of AD_be_- inoculated mice compared to Ctrl_be_-inoculated ones (*p*=0.005, Mann-Whitney test). (**d-f**) Gallyas silver staining revealed labelled neurons evoking NFTs (**e**) as well as Aβ plaques (**f**). *n_Ctrl_*=15, *n_AD1_*=15, *n_AD2_*=20. Data are shown as mean ± s.e.m. Scale bars = 100 µm (**a, d**) and 20 µm (**b, e, f**).

**Supplementary Figure 7.**
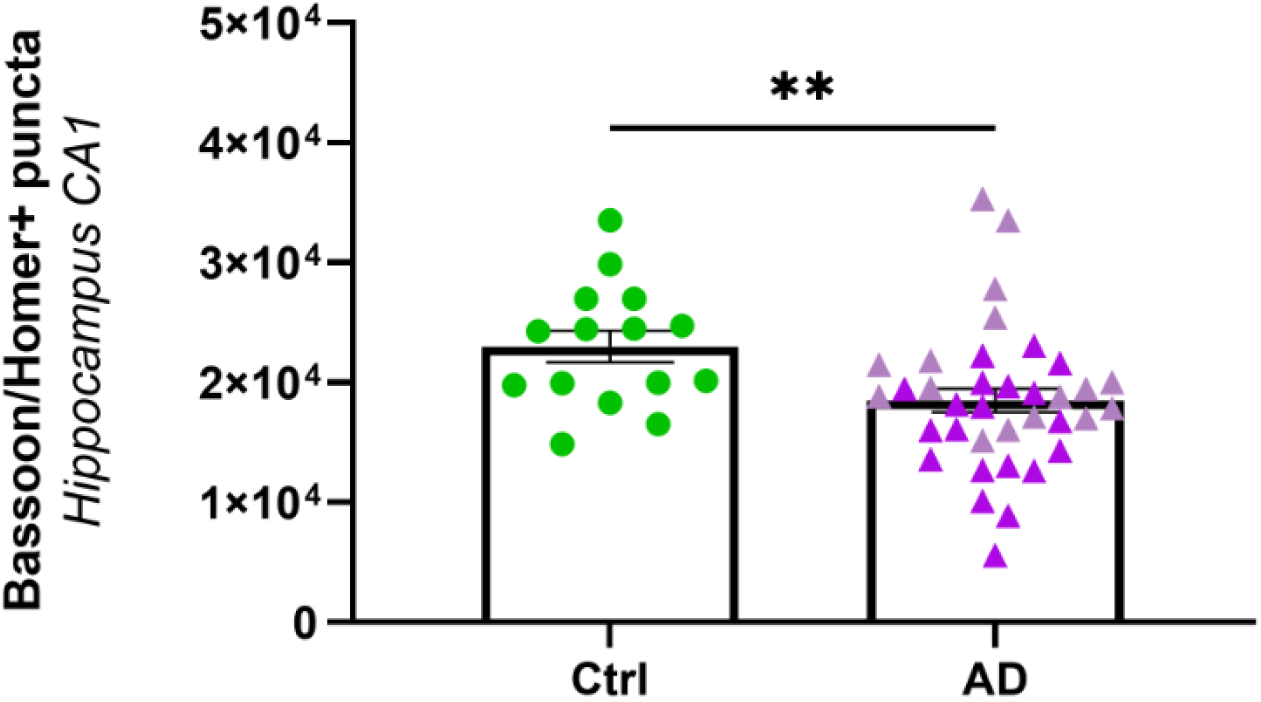
Synaptic density is decreased in the CA1 of AD_be_-inoculated APP_swe_/PS1_dE9_ mice compared to Ctrl_be_-inoculated mice at 8 mpi (Mann Whitney’s test, p=0.005). *n_Ctrl_*=15, *n_AD1_*=15, *n_AD2_*=20 mice. \*\**p*<0.01; Data are shown as mean ± s.e.m.

**Supplementary Figure 8.**
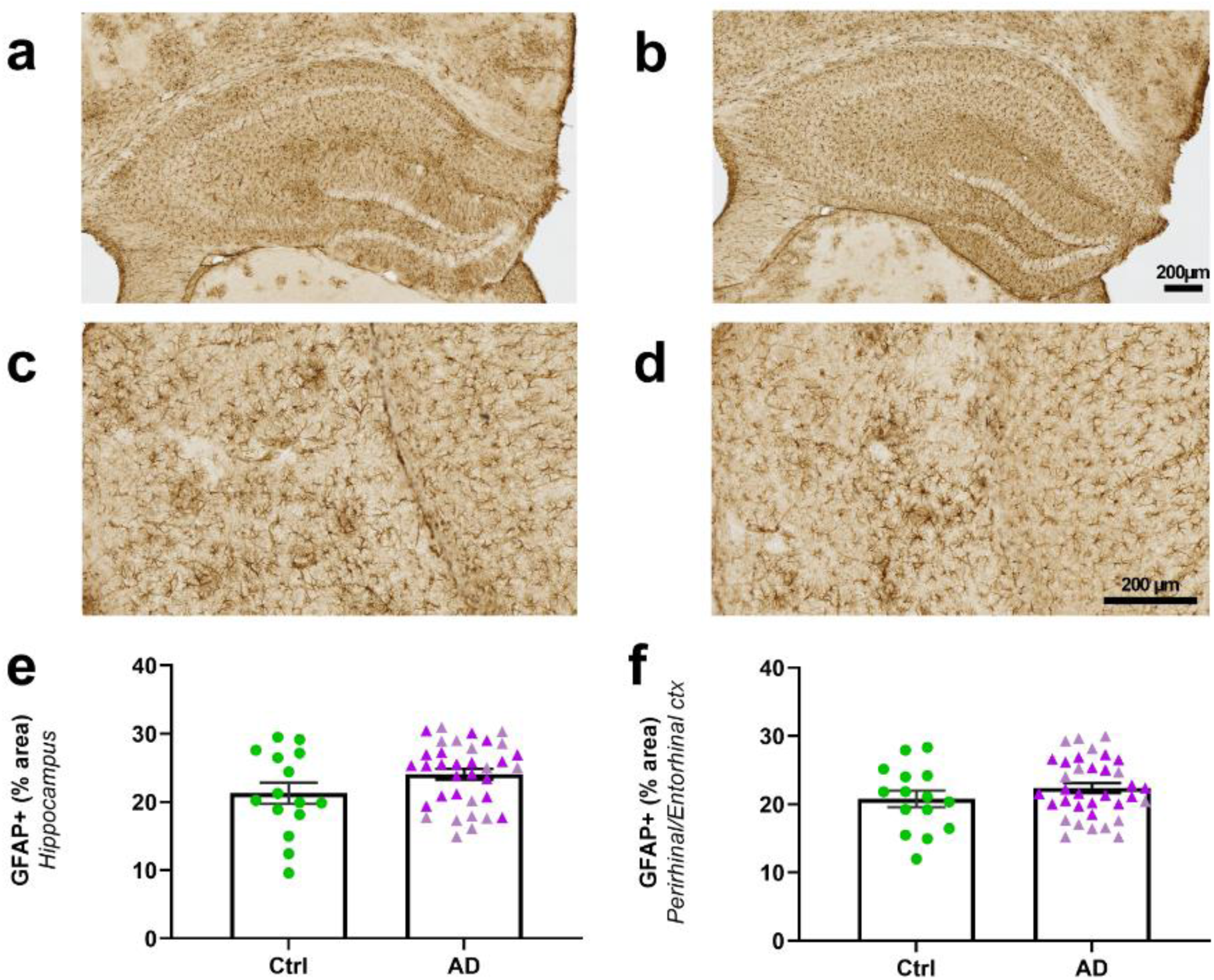
Astrocytic coverage was similar in the hippocampus and perirhinal/entorhinal cortex of AD_be_- and Ctrl_be_-inoculated APP_swe_/PS1_dE9_ mice at 8 mpi. GFAP staining revealed similar astrocytic loads in the hippocampus (**a-b, e**) and perirhinal/entorhinal cortex (**c-d, f**) of AD_be_- and Ctrl_be_-inoculated APP_swe_/PS1_dE9_ mice at 8 mpi. *n_Ctrl_*=15, *n_clAD_*=15, *n_rpAD_*=20 mice. Data are shown as mean ± s.e.m. Scale bars = 200 µm.

**Supplementary Figure 9.**
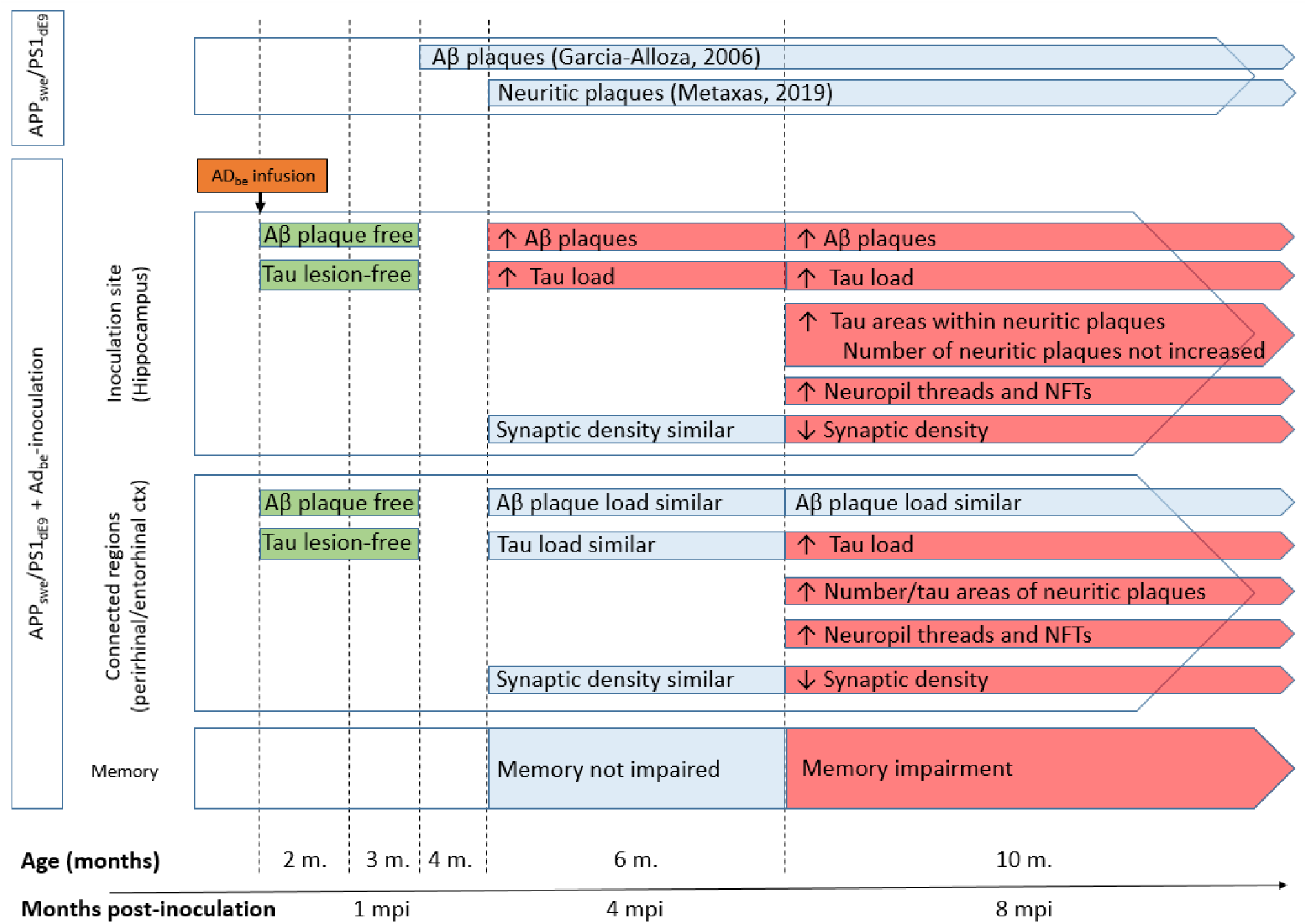
Overview of the changes reported after infusion of AD_be_ in APP_swe_/PS1_dE9_. The first lines (APP_swe_/PS1_dE9_) display published timeframes for occurrence of Aβ plaques (4 month of age) and tau-positive neuritic plaques (6 months of age) in APP_swe_/PS1_dE9_ mice. The lines below (APP_swe_/PS1_dE9_+AD_be_) display events reported in AD_be_- inoculated APP_swe_/PS1_dE9_ mice compared to Ctrl_be_-inoculated APP_swe_/PS1_dE9_ animals. One month post inoculation (mpi), Aβ and tau were not detected in AD_be_-inoculated APP_swe_/PS1_dE9_ mice. Blue frames display events that were similar in AD_be_- and Ctrlbe-inoculated APP_swe_/PS1_dE9_ animals. Red frames highlight impairments detected in AD_be_-inoculated APP_swe_/PS1_dE9_ mice. First changes concerned increased Aβ and tau load within the hippocampus at 4 mpi. Synaptic density and memory were not modified at this stage. Then at 8 mpi, increased Aβ and tau load continued to involve the hippocampus and tau pathology extended to connected regions (perirhinal/entorhinal cortex). Synaptic density was reduced in the hippocampus and perirhinal/entorhinal cortex and memory was impaired at this stage.

## REFERENCES

1 Audouard E, Houben S, Masaracchia C, Yilmaz Z, Suain V, Authelet M, De Decker R, Buee L, Boom A, Leroy K, Ando K, Brion JP (2016) High-molecular-weight paired helical filaments from Alzheimer brain induces seeding of wild-type mouse tau into an argyrophilic 4R Tau pathology in vivo. Am J Pathol 186: 2709–2722. https://doi.org/10.1016/j.ajpath.2016.06.008

2 Bennett DA, Schneider JA, Wilson RS, Bienias JL, Arnold SE (2004) Neurofibrillary tangles mediate the association of amyloid load with clinical Alzheimer disease and level of cognitive function. Arch Neurol 61: 378–384. https://doi.org/10.1001/archneur.61.3.378

3 Braak H, Braak E (1991) Neuropathological stageing of Alzheimer related changes. Acta Neuropathol 82: 239–259. https://doi.org/10.1007/BF00308809

4 Clavaguera F, Bolmont T, Crowther RA, Abramowski D, Frank S, Probst A, Fraser G, Stalder AK, Beibel M, Staufenbiel M, Jucker M, Goedert M, Tolnay M (2009) Transmission and spreading of tauopathy in transgenic mouse brain. Nat Cell Biol 11: 909–913. https://doi.org/10.1038/ncb1901

5 Condello C, Yuan P, Schain A, Grutzendler J (2015) Microglia constitute a barrier that prevents neurotoxic protofibrillar Abeta42 hotspots around plaques. Nat Commun 6: 6176. https://doi.org/10.1038/ncomms7176

6 d’Orange M, Auregan G, Cheramy D, Gaudin-Guerif M, Lieger S, Guillermier M, Stimmer L, Josephine C, Herard AS, Gaillard MC, Petit F, Kiessling MC, Schmitz C, Colin M, Buee L, Panayi F, Diguet E, Brouillet E, Hantraye P, Bemelmans AP, Cambon K (2018) Potentiating tangle formation reduces acute toxicity of soluble tau species in the rat. Brain 41: 535–549. https://doi.org/10.1093/brain/awx342

7 Dejanovic B, Huntley MA, De Maziere A, Meilandt WJ, Wu T, Srinivasan K, Jiang ZY, Gandham V, Friedman BA, Ngu H, Foreman O, Carano RAD, Chih B, Klumperman J, Bakalarski C, Hanson JE, Sheng M (2018) Changes in the synaptic proteome in tauopathy and rescue of Tau-induced synapse loss by C1q antibodies. Neuron 100: 1322–1336.e1327. https://doi.org/10.1016/j.neuron.2018.10.014

8 du Sert NP, Hurst V, Ahluwalia A, Alam S, Avey MT, Baker M, Browne WJ, Clark A, Cuthill IC, Dirnagl U, Emerson M, Garner P, Holgate ST, Howells DW, Karp NA, Lazic SE, Lidster K, MacCallum CJ, Macleod M, Pearl EJ, Petersen OH, Rawle F, Reynolds P, Rooney K, Sena ES, Silberberg SD, Steckler T, Wurbel H (2020) The ARRIVE guidelines 2.0: Updated guidelines for reporting animal research. Bmc Veterinary Research 16: Article 242. https://doi.org/10.1186/s12917-020-02451-y

9 Garcia-Alloza M, Robbins EM, Zhang-Nunes SX, Purcell SM, Betensky RA, Raju S, Prada C, Greenberg SM, Bacskai BJ, Frosch MP (2006) Characterization of amyloid deposition in the APPswe/PS1dE9 mouse model of Alzheimer disease. Neurobiol Dis 24: 516–524. https://doi.org/10.1016/j.nbd.2006.08.017

10 Gary C, Lam S, Herard AS, Koch JE, Petit F, Gipchtein P, Sawiak SJ, Caillierez R, Eddarkaoui S, Colin M, Aujard F, Deslys JP, French Neuropathology Network, Brouillet E, Buée L, Comoy EE, Pifferi F, Picq J-L, Dhenain M (2019) Encephalopathy induced by Alzheimer brain inoculation in a non-human primate. Acta Neuropathol Commun 7: 126. https://doi.org/10.1186/s40478-019-0771-x

11 Gilles JF, Dos Santos M, Boudier T, Bolte S, Heck N (2017) DiAna, an ImageJ tool for object-based 3D co-localization and distance analysis. Methods 115: 55–64. https://doi.org/10.1016/j.ymeth.2016.11.016

12 Guo JL, Narasimhan S, Changolkar L, He ZH, Stieber A, Zhang B, Gathagan RJ, Iba M, McBride JD, Trojanowski JQ, Lee VMY (2016) Unique pathological tau conformers from Alzheimer’s brains transmit tau pathology in nontransgenic mice. J Exp Med 213: 2635–2654. https://doi.org/10.1084/jem.20160833

13 He ZH, Guo JL, McBride JD, Narasimhan S, Kim H, Changolkar L, Zhang B, Gathagan RJ, Yue CY, Dengler C, Stieber A, Nitla M, Coulter DA, Abel T, Brunden KR, Trojanowski JQ, Lee VMY (2018) Amyloid-beta plaques enhance Alzheimer’s brain tau-seeded pathologies by facilitating neuritic plaque tau aggregation. Nat Med 24: 29–38. https://doi.org/10.1038/nm.4443

14 Hoover BR, Reed MN, Su J, Penrod RD, Kotilinek LA, Grant MK, Pitstick R, Carlson GA, Lanier LM, Yuan LL, Ashe KH, Liao D (2010) Tau mislocalization to dendritic spines mediates synaptic dysfunction independently of neurodegeneration. Neuron 68: 1067–1081. https://doi.org/10.1016/j.neuron.2010.11.030

15 Lam S, Petit F, Hérard A-S, Boluda S, Eddarkaoui S, Guillermier M, The Brainbank Neuro-CEB Neuropathology Network, Buée L, Duyckaerts C, Haïk S, Picq J-L, Dhenain M (2021) Transmission of amyloid-beta and tau pathologies is associated with cognitive impairments in a primate. Acta Neuropathol Commun 9: 165. https://doi.org/10.1186/s40478-021-01266-8

16 Leyns CEG, Gratuze M, Narasimhan S, Jain N, Koscal LJ, Jiang H, Manis M, Colonna M, Lee VMY, Ulrich JD, Holtzman DM (2019) TREM2 function impedes tau seeding in neuritic plaques. Nat Neurosci 22: 1217–1222. https://doi.org/10.1038/s41593-019-0433-0

17 Metaxas A, Thygesen C, Kempf SJ, Anzalone M, Vaitheeswaran R, Petersen S, Landau AM, Audrain H, Teeling JL, Darvesh S, Brooks DJ, Larsen MR, Finsen B (2019) Ageing and amyloidosis underlie the molecular and pathological alterations of tau in a mouse model of familial Alzheimer’s disease. Sci Rep 9: Article number 15758. https://doi.org/10.1038/S41598-019-52357-5

18 Nelson PT, Alafuzoff I, Bigio EH, Bouras C, Braak H, Cairns NJ, Castellani RJ, Crain BJ, Davies P, Del Tredici K, Duyckaerts C, Frosch MP, Haroutunian V, Hof PR, Hulette CM, Hyman BT, Iwatsubo T, Jellinger KA, Jicha GA, Kovari E, Kukull WA, Leverenz JB, Love S, Mackenzie IR, Mann DM, Masliah E, McKee AC, Montine TJ, Morris JC, Schneider JA, Sonnen JA, Thal DR, Trojanowski JQ, Troncoso JC, Wisniewski T, Woltjer RL, Beach TG (2012) Correlation of Alzheimer disease neuropathologic changes with cognitive status: a review of the literature. J Neuropath Exp Neur 71: 362–381. https://doi.org/10.1097/NEN.0b013e31825018f7

19 Noda-Saita K, Terai K, Iwai A, Tsukamoto M, Shitaka Y, Kawabata S, Okada M, Yamaguchi T (2004) Exclusive association and simultaneous appearance of congophilic plaques and AT8-positive dystrophic neurites in Tg2576 mice suggest a mechanism of senile plaque formation and progression of neuritic dystrophy in Alzheimer’s disease. Acta Neuropathol 108: 435–442. https://doi.org/10.1007/s00401-004-0907-2

20 Olmos-Alonso A, Schetters STT, Sri S, Askew K, Mancuso R, Vargas-Caballero M, Holscher C, Perry VH, Gomez-Nicola D (2016) Pharmacological targeting of CSF1R inhibits microglial proliferation and prevents the progression of Alzheimer’s-like pathology. Brain 139: 891–907. https://doi.org/10.1093/brain/awv379

21 Paxinos G, Franklin KBJ (2001) The mouse brain in stereotaxic coordinates. Academic Press, City

22 Pooler AM, Noble W, Hanger DP (2014) A role for tau at the synapse in Alzheimer’s disease pathogenesis. Neuropharmacology 76: 1–8. https://doi.org/10.1016/j.neuropharm.2013.09.018

23 Radde R, Bolmont T, Kaeser SA, Coomaraswamy J, Lindau D, Stoltze L, Calhoun ME, Jaggi F, Wolburg H, Gengler S, Haass C, Ghetti B, Czech C, Holscher C, Mathews PM, Jucker M (2006) Abeta42-driven cerebral amyloidosis in transgenic mice reveals early and robust pathology. EMBO Rep 7: 940–946. https://doi.org/10.1038/sj.embor.7400784

24 Roberson ED, Scearce-Levie K, Palop JJ, Yan F, Cheng IH, Wu T, Gerstein H, Yu GQ, Mucke L (2007) Reducing endogenous tau ameliorates amyloid beta-induced deficits in an Alzheimer’s disease mouse model. Science 316: 750–754. https://doi.org/10.1126/science.1141736

25 Sanchez-Mejias E, Navarro V, Jimenez S, Sanchez-Mico M, Sanchez-Varo R, Nunez-Diaz C, Trujillo-Estrada L, Davila JC, Vizuete M, Gutierrez A, Vitorica J (2016) Soluble phospho-tau from Alzheimer’s disease hippocampus drives microglial degeneration. Acta Neuropathol 132: 897–916. https://doi.org/10.1007/s00401-016-1630-5

26 Selkoe DJ (2002) Alzheimer’s disease is a synaptic failure. Science 298: 789–791. https://doi.org/10.1126/science.1074069

27 Sivanesan S, Tan A, Rajadas J (2013) Pathogenesis of abeta oligomers in synaptic failure. Curr Alzh Res 10: 316–323. https://doi.org/10.2174/1567205011310030011

28 Spires-Jones TL, Hyman BT (2014) The intersection of amyloid beta and tau at synapses in Alzheimer’s disease. Neuron 82: 756–771. https://doi.org/10.1016/j.neuron.2014.05.004

29 Streit WJ, Braak H, Xue QS, Bechmann I (2009) Dystrophic (senescent) rather than activated microglial cells are associated with tau pathology and likely precede neurodegeneration in Alzheimer’s disease. Acta Neuropathol 118: 475–485. https://doi.org/10.1007/s00401-009-0556-6

30 Terry RD, Masliah E, Salmon DP, Butters N, DeTeresa R, Hill R, Hansen LA, Katzman R (1991) Physical basis of cognitive alterations in Alzheimer’s disease: synapse loss is the major correlate of cognitive impairment. Ann Neurol 30: 572–580. https://doi.org/10.1002/ana.410300410

31 Vergara C, Houben S, Suain V, Yilmaz Z, De Decker R, Vanden Dries V, Boom A, Mansour S, Leroy K, Ando K, Brion JP (2019) Amyloid-pathology enhances pathological fibrillary tau seeding induced by Alzheimer PHF in vivo. Acta Neuropathol 137: 397–412. https://doi.org/10.1007/s00401-018-1953-5

